# Psychophysical reverse correlation reflects both sensory and decision-making processes

**DOI:** 10.1101/273680

**Authors:** Gouki Okazawa, Long Sha, Braden A. Purcell, Roozbeh Kiani

**Author notes:** **Corresponding Author:** Roozbeh Kiani, Center for Neural Science, New York University, 4 Washington Pl, Room 809, New York, NY 10003.

## Abstract

Goal directed behavior depends on both sensory mechanisms that gather information from the outside world and decision-making mechanisms that select appropriate behavior based on that sensory information. Psychophysical reverse correlation is commonly used to quantify how fluctuations of sensory stimuli influence behavior and is generally believed to uncover the spatiotemporal weighting functions of sensory processes. Here we show that reverse correlations also reflect decision-making processes and can deviate significantly from the true sensory filters. Specifically, changes of decision bound and mechanisms of evidence integration systematically alter psychophysical reverse correlations. Similarly, trial-to-trial variability of sensory and motor delays and decision times causes systematic distortions in psychophysical kernels that should not be attributed to sensory mechanisms. We show that ignoring details of the decision-making process results in misinterpretation of reverse correlations, but proper use of these details turns reverse correlation into a powerful method for studying both sensory and decision-making mechanisms.

## Introduction

Accurate characterization of behavior is key to understanding neural computations^1,2^. Not only do we want to know which behaviors arise from sensory inputs in an environment, but also we need to understand the mechanisms through which sensory inputs lead to behavioral outputs. Over the past decades, several system identification techniques have been developed to address these needs. Among the most commonly used is psychophysical reverse correlation^3–5^, a technique that aims to estimate how sensory information is weighted to guide decisions. The core idea is that by quantifying the stimulus fluctuations that precede each choice (i.e., reverse correlation), one can infer the spatiotemporal filter implemented by the sensory processes (Fig. 1). It can be shown mathematically that under the assumptions of signal detection theory (SDT) for the decision-making process, psychophysical reverse correlation does recover the true sensory weights^6,7^. In SDT, a linear filter is applied to a sensory stimulus and the outcome is compared to a decision criterion. The result of this comparison (higher or lower than the criterion) dictates the choice^8,9^. If stimuli on different trials are drawn from a symmetric distribution (e.g., Gaussian) reverse correlation will accurately estimate the linear sensory filter of SDT by averaging the stimuli that precede a particular choice.

**Figure 1.**
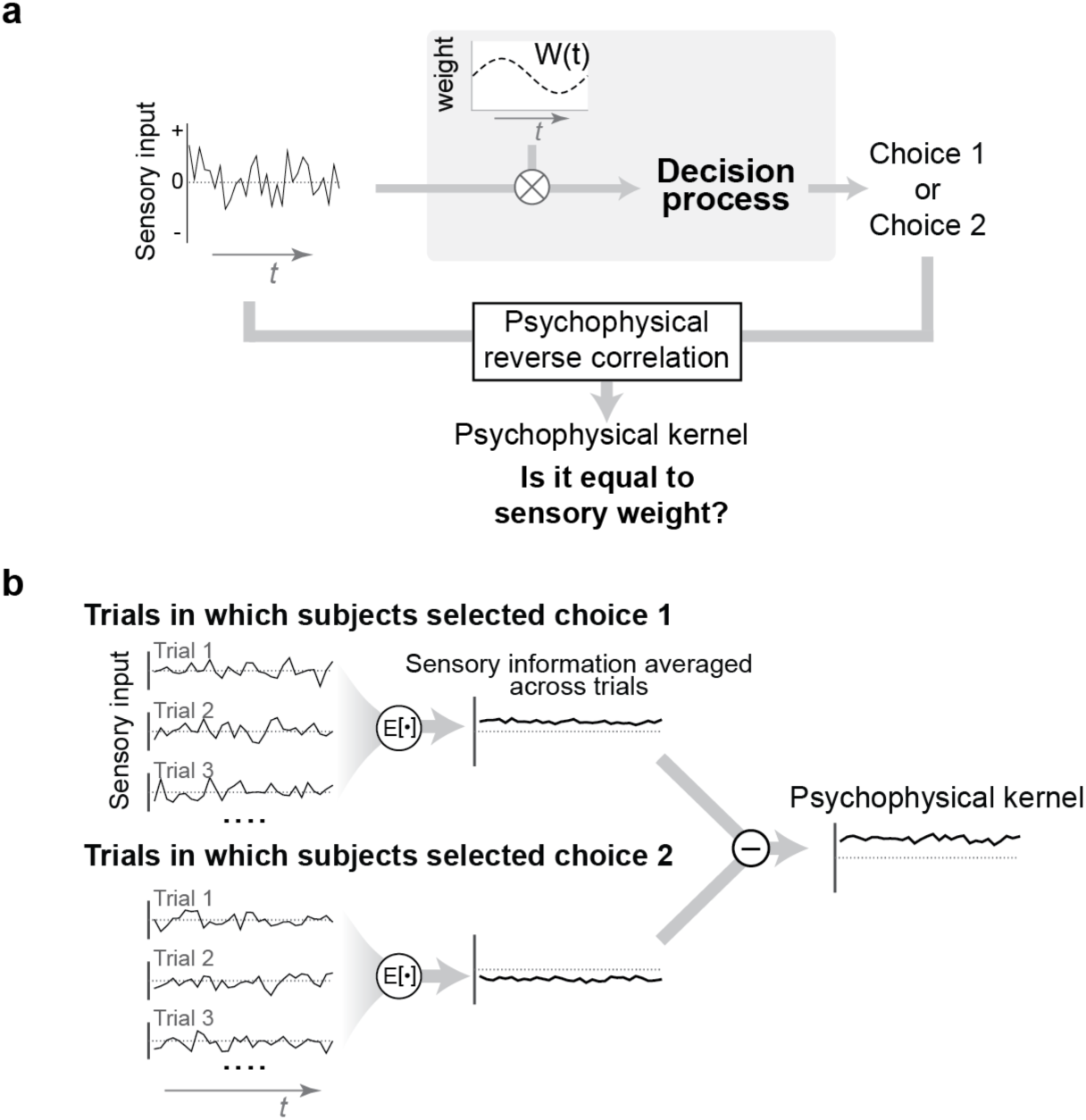
Psychophysical reverse correlation has been developed to recover sensory weights in perceptual tasks but it could also be influenced by decision-making mechanisms. (**a**) In a typical reverse correlation experiment, subjects receive a sequence of randomly fluctuating sensory information and make a binary choice. Experimenters can directly observe the stimulus and choice, but not the sensory weights and decision-making process (gray box). When choices are made by applying a sensory filter (weighting function) to the stimulus and comparing the result against a criterion, as proposed by signal detection theory, psychophysical reverse correlation will recover the sensory filter. However, it is unknown how well the analysis generalizes to more complex decision-making mechanisms. (**b**) Reverse correlation calculates the average stimuli preceding each choice and subtracts the results for the two choices. The outcome is a “psychophysical kernel.”

The technique can also be extended to the temporal domain to recover the dynamics of the weighting function when choices are based on filtering a sequence of observations and comparing the results to a criterion^5,10^. These temporal extensions resemble spike-triggered averaging techniques, which derive spatiotemporal receptive fields (linear kernels) of spiking neurons^11–16^, under the assumption that firing rates are determined by filtering sensory inputs followed by application of a static nonlinearity. In general, when a discrete outcome arises from a sequence of linear and nonlinear computations, reverse correlation is a recommended method for estimating the linear component of the computation. How well does this recommendation work in practice for sensory decisions?

Studies of the decision-making process over the past decade have revealed that the simple assumptions of SDT do not adequately capture the complexity of perceptual decisions. We now know that for many decisions, subjects integrate sensory evidence in favor of different choices, and the final decision is made when the integrated evidence reaches a satisfactory threshold^17–22^. Several key features of this process are absent in simple temporal extensions of SDT. First, subjects can flexibly adjust their decision bound within and across trials to change how much evidence to integrate, and thereby trade off accuracy and speed of their decisions^23,24^. Second, neural implementation of the decision-making process relies on a competition or race between multiple integrators, rather than reaching a decision bound in a single integrator. Third, realistic implementations of these computations in neural networks require taking into account biophysical constraints (e.g., lower limit of firing rates at zero^25,26,27^) and network mechanisms of integration (e.g., mutual inhibition^21,22,25^). Finally, applying theory to real experimental data requires taking practical limitations into account. A key factor that has been largely ignored thus far is the sensory and motor delays (non-decision time). The sum of the non-decision time and the time spent on integration of evidence (decision time) determine experimentally measured reaction times (RTs)^28–30^. Because the non-decision time limits the relevant stimulus history for the choice, it could distort the outcome of reverse correlation. How much do these factors influence the estimation of sensory filters with psychophysical reverse correlation? Except for scant examples in the past literature that studied basic properties of the integration of sensory evidence (e.g., bounded or leaky accumulation)^25,31–33^, the answer is largely unknown. A systematic exploration is timely because mechanistic studies of sensory and decision-making processes have become a cornerstone of modern neuroscience and because accurate methods for quantifying the relationship between experimental stimuli and behavior provide a critical foundation for these investigations^2^.

We show that psychophysical reverse correlation deviates qualitatively and quantitatively from sensory weights under several variants of decision-making models. Experiments in which stimulus viewing duration is controlled by the experimenter often do not allow distinguishing these variants, leaving the mechanistic cause of observed kernel dynamics obscure, unless special measures are implemented (e.g., variation of stimulus durations across trials). RT tasks, where the stimulus-viewing duration is controlled by the subject and reaction times can be directly measured by experimenters, offer much more leverage, especially when a model-based approach is adopted to correct for expected deviations of the reverse correlation from sensory weights. We show that these deviations are not caused by the presence of a decision bound. Rather, they emerge from the presence of variable sensory and motor delays, changes of decision-bound within and across trials, lower limits for accumulated evidence, integration time constants, and mutual inhibition of competing accumulators. Knowing about these deviations enables us to correct for them, when possible, and prevents false conclusions about temporal variation of sensory weights. We demonstrate this point in a series of experiments by showing that the mechanism that underlies decisions predicts temporal dynamics of psychophysical kernels and quantitatively explains experimentally derived kernels.

## Results

In a typical reverse correlation experiment, subjects observe a sequence of noisy sensory stimuli and try to detect the presence of a target or categorize a stimulus^3,4,32–35^ (Fig. 1). The stimuli could be a random dot kinematogram^31,32^, oriented gratings or bars^5,36^, or any other sensory inputs that randomly vary within or across trials along one or more stimulus attributes. For example, in the random-dot direction discrimination task, motion energy for a 0% coherence stimulus fluctuates from moment to moment according to a bell-shaped distribution centered on zero^32^ (positive and negative values correspond to net motion in the two discriminated directions). The reverse correlation analysis calculates the relationship between subjects’ choice and moment-to-moment stimulus fluctuations by averaging over the stimuli that precede a particular choice. A common observation is that when there happens to be more rightward motion in a trial, subjects are more likely to choose right and vice versa^31,32,37^. For two-alternative decision tasks, the analysis yields two kernels, one for each choice. Because of symmetry of the two choices, the kernels tend to be mirror images of each other^32,38^. Therefore, it is customary to subtract the two kernels and report the result (Fig. 1b):

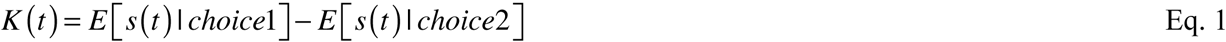

where *E*[*s*(*t*)|*choice*1] indicates the trial average of the stimulus at time *t* conditional on choice 1, *s*(*t*) is the stimulus drawn from a stochastic function with symmetric noise (e.g., Gaussian), and *K*(*t*) is the magnitude of the psychophysical kernel at time *t*.

Psychophysical kernels are guaranteed to match the sensory filters when decisions are made by applying a static nonlinearity^6,7,11,39^, for example, comparison to a decision criterion, as suggested by SDT^8,9^. However, recent advances suggest that SDT offers an incomplete characterization of the decision-making process. In particular, many perceptual decisions depend on integration of sensory information toward a decision bound^17–20,25,33,40,41^, the decision bound can vary based on speed-accuracy tradeoff^23,24^, the integration is influenced by urgency^42–44^ and prior signals^40,45–47^, and experimentally measured RTs consist of a combination of decision times and non-decision times^28–30^.

A simple and commonly used class of decision-making models that takes these intricacies into account and provides a quantitative explanation of behavior in perceptual tasks is the drift diffusion model (DDM)^17,18,48^ and its extensions^24,26,49,50^. In these models, weighted sensory evidence is integrated over time until the integrated evidence (the decision variable, DV) reaches either an upper (positive) or a lower (negative) bound (Fig. 2), where each bound corresponds to one of the choices. We begin our exploration with the most basic model but will focus on more complex implementations later in the paper. Our conclusions in this section are not limited to a specific implementation and generalize to a wide variety of models in this class.

**Figure 2.**
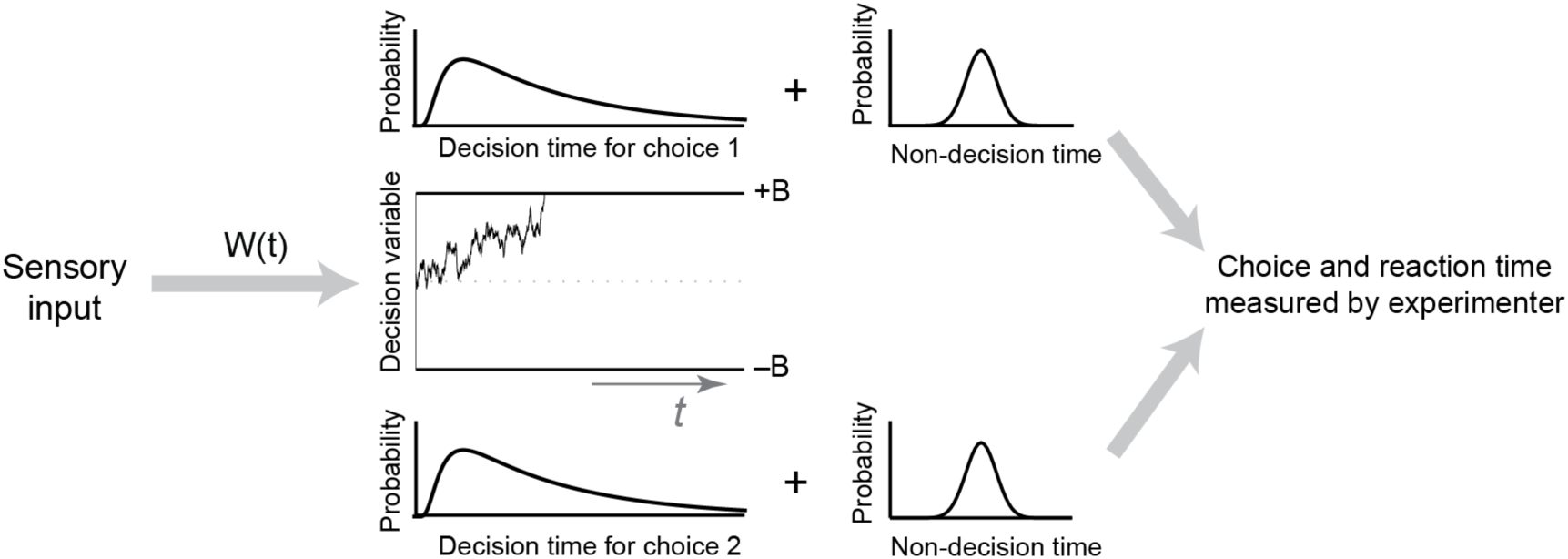
The drift diffusion model (DDM) captures the core computations for perceptual decisions made by integration of sensory information over time. We use variants of this model and more sophisticated extensions to explore how the decision-making mechanism influences psychophysical kernels. In DDMs, a weighting function, *w*(*t*), is applied to the sensory inputs to generate the momentary evidence, which is integrated over time to form the decision variable (DV). The DV fluctuates over time due to changes in the sensory stimulus and neural noise for stimulus representation and integration. As soon as the DV reaches one of the two decision bounds (+*B* for choice 1 and –*B* for choice 2), the integration terminates and a choice is made (decision time). However, reporting the choice happens after a temporal gap due to sensory and motor delays (non-decision time). Experimenters know about the choice after this gap and can measure only the reaction time (the sum of decision and non-decision times) but not the decision time.

Neither the integration process nor the boundedness of the integration per se causes a systematic deviation between the psychophysical kernels and true sensory weights. We define true sensory weights as the weights applied to the sensory information to create the momentary evidence that will be accumulated over time for making a decision. Psychophysical kernels are the best approximation of these weights under the assumptions of SDT. We explore how close this match is under DDM. In Methods, we provide a mathematical proof that in a plain version of the DDM where decision bound and noise are constant over time and behavioral responses are generated as soon as the DV reaches one of the bounds (non-decision time = 0), psychophysical kernels are proportional to the sensory weights:

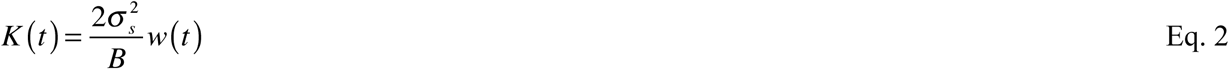

where *w*(*t*) is the time-dependent weight applied to sensory inputs prior to integration, 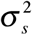 is the variance of stimulus fluctuations, and *B* is the height of the decision bound. Similar results can be obtained for unbounded DDMs (Eq. 14, see Methods). Fig. 3 shows simulations that confirm our proofs. Reverse correlation for an unbounded integration process with constant or sinusoidally varying weights recovers the true weighting function (Fig. 3a-c, S1). Similarly, it yields the true weights for a bounded DDM, (Fig. 3d-e, h), regardless of the decision bound height.

**Figure 3.**
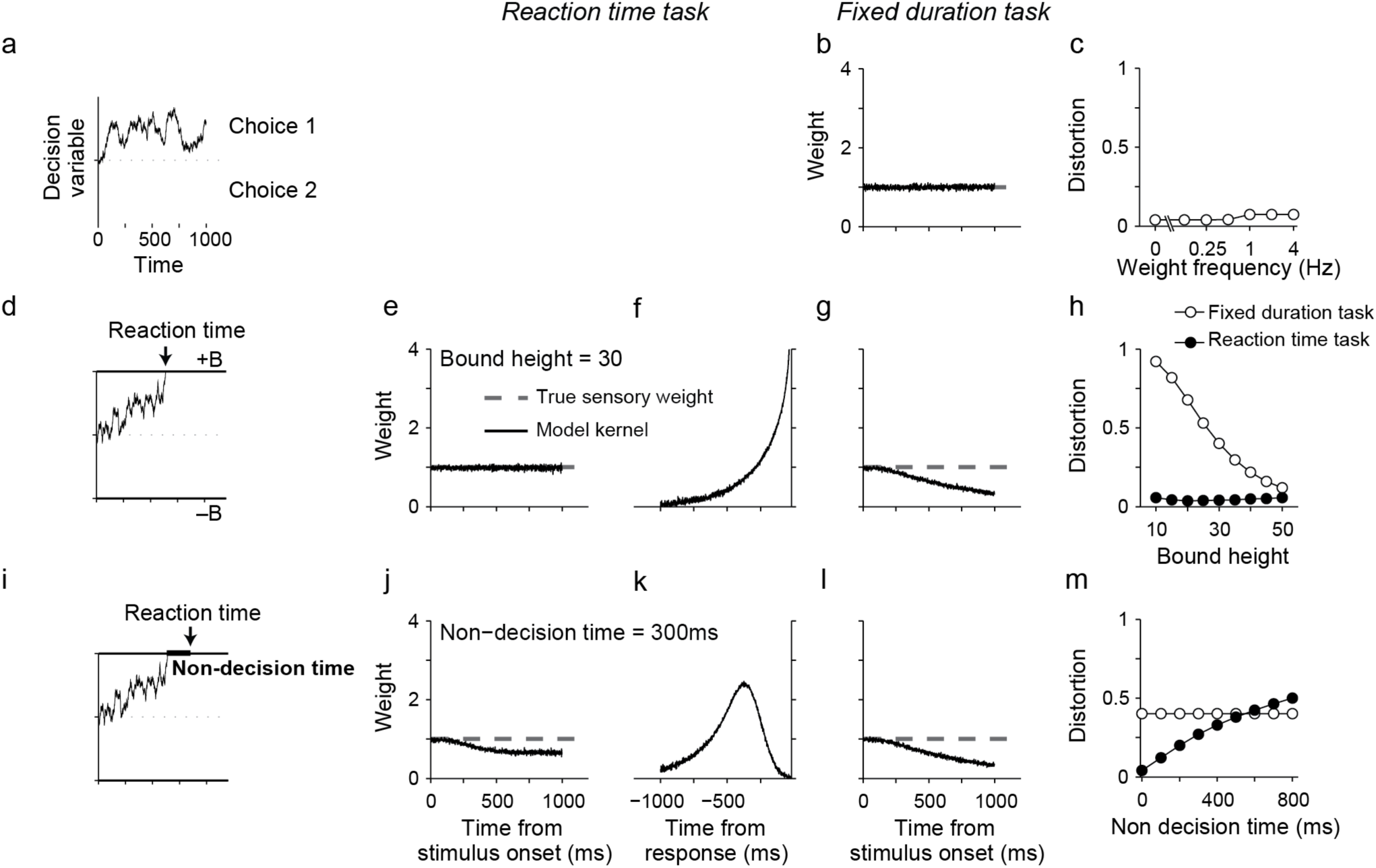
Psychophysical kernels deviate from sensory weights in DDM because of incomplete knowledge about decision time. (**a-c**) Integration of evidence per se does not preclude accurate recovery of sensory weights. For an unbounded DDM that integrates the momentary evidence as long as sensory inputs are available, the psychophysical kernel matches the true sensory weights. In this simulation, the weight is stationary and fixed at 1, but similarly matching results are obtained for any sensory weight (c). Distortion quantifies root mean square error between the psychophysical kernel and the true sensory weights (Eq. 17). See Fig. S1 for additional examples. Unbounded DDMs are unrealistic for RT tasks because they lack a termination criterion. (**d-h**) The decision bound does not preclude accurate recovery of sensory weights. In a bounded DDM without non-decision time, RTs are identical to decision times (d). Model simulations for RT tasks result in stimulus-aligned kernels that match sensory weights of the model (e) and response-aligned kernels that rise monotonically (f), as expected for termination with bound crossing. However, stimulus-aligned kernels in fixed-duration tasks show a monotonic decrease because later stimuli are less likely to bear on the choice (g). This deviation from sensory weights gets smaller as the decision bound rises and commitment to a choice before termination of the stimulus becomes less likely (h). (**i-m**) Variability of nondecision time makes reaction time an unreliable estimate of decision time, causing systematic deviations between psychophysical kernels and true sensory weights. After including non-decision time in the bounded DDM, stimulus-aligned kernel in RT tasks show a monotonic decrease because stimuli that immediately precede the choice do not contribute to it (j). Response-aligned kernels show a peak, whose time is dependent on the distribution of non-decision times (k). Kernels for fixed-duration tasks are not affected by non-decision time, if there is a long enough delay between the stimulus onset and response cue (l). However, they still show the decline caused by bound crossing, similar to g. Deviation of stimulus-aligned kernels in the RT task increases with variability of non-decision time (m). Standard deviation of non-decision time is assumed to be 1/3 of its mean in these simulations. All kernels are normalized according to Eq. 2 or Eq. 14 to allow direct comparison with the true sensory weights (see Methods).

Although the proportionality in Eq. 2 may suggest that psychophysical kernels can be successfully used to recover spatiotemporal dynamics of sensory weights, critical limitations prevent that in practice, as we explain below. The most common limitation is experimenter’s lack of knowledge about decision time, which is caused by asynchrony between the time that the DV reaches a decision bound (bound-crossing time or decision time) and the subject’s report of the decision (when the choice becomes known to the experimenter). Such asynchronies stem from two sources: delays in neural circuitry and experimental design.

In many experiments subjects are exposed to the stimulus for a duration determined by the experimenter and can report their choice only after a Go cue. In these “fixed-duration” designs, the exact decision time and its trial-to-trial variability are unknown to the experimenter, and decision times are likely to be prior to the Go cue^32,51^. Because stimuli presented after the bound-crossing time do not contribute to the choice (or contribute less)^31,52,53^, including that period in the calculation of psychophysical kernels leads to a progressive underestimation of sensory weights^31,32,54^, causing a systematic deviation from Eq. 2 (Fig. 3g-h), compatible with past studies^55^. The diminishing kernel (Fig. 3g) correctly characterizes the effective reduction of the influence of the sensory stimulus on choice. However, note that from an experimenter’s perspective, the shape of the kernel is inadequate to tell whether the reduced influence of the stimulus on choice is caused by a change in sensory weights, by early termination of the decision during stimulus viewing, by a combination of both, or by another mechanism in the decision-making process (see below). Such a mechanistic understanding could be achieved only if the experimental design is enriched and a model-based approach is adopted. Although there are successful examples of achieving such goals^31,32^, fixed-duration tasks impose significant limitations on experimenters’ ability to determine the beginning and end of the decision-making process (cf. ref. 51), which would be necessary for separating sensory and decision-making mechanisms that shape psychophysical kernels.

Experimental designs in which subjects respond as soon as they make their decision (RT tasks; Fig. 3e) enable measurement of decision times and can be used to address the problem. However, RT tasks come with their own challenges for psychophysical reverse correlations. Sensory and motor delays are among them (Fig. 3i). Although the presence of such delays is widely appreciated, their effect on psychophysical kernels is unexplored. These delays effectively create a temporal gap between bound crossing and the report of the decision, making stimuli immediately before the report inconsequential for the decision. Fig. 3j shows that non-decision times pull down the psychophysical kernel. These systematic reductions can cause the illusion of non-stationarity for stationary sensory weights (Fig. 3j, m) or distort the dynamics of time-varying weights (Fig. S2).

What makes the psychophysical reverse correlation especially vulnerable to non-decision times is the variable nature of the sensory and motor delays^56–58^. A fixed non-decision time would cause a readily detectable signature (Fig. S3) and is easy to correct for by excluding the last stimulus fluctuations in each trial that corresponded to the non-decision time. Similarly, if the non-decision time was variable but we could know the exact delay on each trial we could easily discard the corresponding period at the end of the stimulus before calculating the kernels to correct for the artificial dynamics caused by the non-decision time. In practice, the non-decision time is not a fixed number^30^. Further, the variability of non-decision time is often in the same order of magnitude as the decision time^26,41,59–61^, making it challenging to thoroughly scrub away the effect of non-decision time just by trimming the stimuli. A more efficient solution is to embrace the distortion caused by the non-decision time, develop an explicit model of both the sensory and decision-making mechanisms, and compare the predictions of such a model with experimentally derived kernels (see the next section).

The fixed-duration design is not affected by the non-decision time, if there is a long enough delay between the stimulus and Go cue or if the stimulus duration is long enough to exceed the tail of the reaction time distribution in an equivalent RT task design (Fig. 3l-m). However, as mentioned above, lack of knowledge about the beginning and end of the integration process in fixed-duration tasks impedes mechanistic studies of kernel dynamics.

So far, we have focused on psychophysical kernels aligned to the stimulus onset. In a RT task, the stimulus viewing duration varies from trial to trial and we can choose to align the kernel to subjects’ responses. Such an alignment is informative both about the termination mechanism of the decision-making process and about the distribution of non-decision times. When the decision-making process stops by reaching a decision bound, the kernel is guaranteed to show a steep rise close to the decision time (Fig. 3f) because stopping is conditional on a stimulus fluctuation that takes the DV beyond the bound. This rise of the kernel does not indicate an increase of sensory weights immediately before the decision. Further, the magnitude of this rise is not always fixed and depends on the decision bound and distribution of non-decision times (see below; Fig. S3). In the presence of a variable non-decision time (Fig. 3k), response-aligned kernels peak and then drop down to zero before the response. The drop happens because the non-decision time causes later fluctuations in the stimulus not to bear on the choice^52,60,62^. The difference between the peak of the kernel and the reaction time is dependent on the mean and standard deviation of the non-decision time, as well as the skewness of its distribution (Fig. S3). Since it is known that the distribution of non-decision times can be quite diverse, depending on the experimental design^63^, the shape of the response-aligned psychophysical kernels can provide an important clue about the distribution of non-decision times and also verification of model-based attempts to discover the non-decision time distribution^63^. Overall, psychophysical kernels aligned to the response are influenced by sensory weights, termination criterion of the decision, and the non-decision time. Consequently, they reflect both sensory and decision-making mechanisms.

## Experimentally measured psychophysical kernels confirm model predictions

The results of the previous section suggest that psychophysical kernels reflect a mixture of sensory and decision-making processes. By embracing this complexity, one can leverage psychophysical kernels to gain insight about both processes. The key is to develop explicit models for both and to compare model predictions against experimentally derived kernels. Below we highlight two experiments designed to achieve this goal.

The first experiment is an RT version of the direction discrimination task^26,64,65^. On each trial, subjects viewed a random dot stimulus and made a saccadic eye movement to one of the two targets as soon as they were ready to report their choice (Fig. 4a). Consistent with previous studies, subjects’ accuracy improved and RTs decreased monotonically with motion strength (Fig. 4b-c)^26,64,65^. We quantified moment-to-moment fluctuations of motion in each trial by calculating the motion energy^32,66,67^ (see Methods). Fig. 4d shows the average and standard deviation of motion energies across all 0% coherence trials (solid black line) and four single-trial examples (dashed black lines). As expected, single trial motion energies departed from 0 with a short latency^66^ (see Methods) and then fluctuated between positive and negative values, which corresponded to the two motion directions discriminated by subjects. Across all 0% coherence trials, these fluctuations canceled each other out, resulting in a zero mean (solid black line) but the standard deviation (gray shading) remained large, indicating short bouts of varying motion strengths in either direction throughout the trial (see Fig. S4 for motion energies of other coherences). The stochastic nature of the stimulus and the known effect of motion energy on the choice^32,52,67^ provided an excellent opportunity to quantify how stimulus dynamics shaped the behavior.

**Figure 4.**
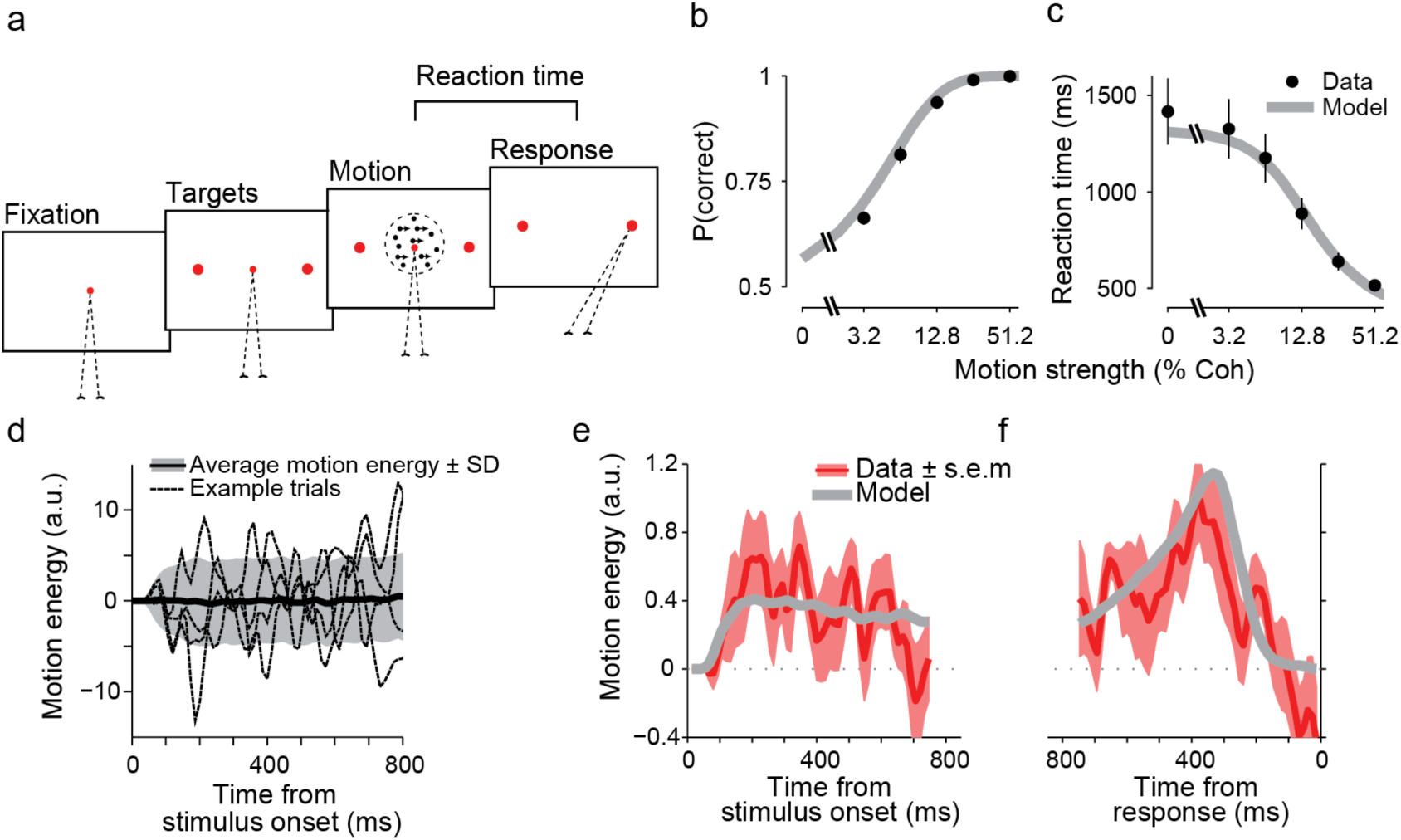
Psychophysical kernels in the direction discrimination task match predictions of a bounded DDM with non-decision time. (**a**) RT task design. Subjects initiated each trial by fixating on a central fixation point. Two targets appeared after a short delay, followed by the random dots stimulus. When ready, subjects indicated their perceived motion direction with a saccadic eye movement to a choice target. The net motion strength (coherence) varied from trial to trial, but also fluctuated within trials due to the stochastic nature of the stimulus. (**b-c**) Choice accuracy increased and RTs decreased with motion strength. Data points are averages across thirteen subjects. Accuracy for 0% motion coherence is 0.5 by design and therefore not shown. Gray lines are fits of a bounded DDM with non-decision time. Error bars denote s.e.m across subjects. (**d**) Motion energy of example 0% coherence trials (dotted lines), and the average (solid black line) and standard deviation (shading) of motion energy across all 0% coherence trials. Positive and negative motion energies indicate the two opposite motion directions in the task. (**e-f**) The bounded DDM predicts psychophysical kernels (gray lines), which accurately match the dynamics of subjects’ kernels (red lines). Because the model sensory weights are stationary, kernel dynamics in the model are caused by the decision-making process and non-decision times. Kernels are calculated for 0% coherence trials. Shading indicates s.e.m across subjects. All kernels are shown up to the minimum of the median RTs across subjects.

We calculated psychophysical kernels using motion energy fluctuations of the 0% coherence trials. Experimentally derived kernels aligned to the stimulus onset or saccade onset (Fig. 4e-f, red lines) were not constant over time and showed a clear non-stationarity. However, their dynamics bore remarkable resemblance to the kernels expected from a DDM with non-decision time and stationary sensory weights (Fig. 3j-k). The main notable difference was a delayed rise in the psychophysical kernel aligned to the stimulus onset (Fig. 4e). As explained above, this delay was inherent to the motion energy calculation, as shown in Fig. 4d.

The qualitative match between experimental and predicted model kernels based on simulations supported the hypothesis that the dynamics in the psychophysical kernels could indeed reflect characteristics of the decision-making process (bound crossing and non-decision time), rather than time-varying sensory weights. We quantitatively tested this hypothesis by fitting the DDM to subjects’ choices and RTs and generating a model prediction for the psychophysical kernels (see Methods). Consistent with past studies, the distribution of RTs and choices across trials provided adequate constraints for estimating all model parameters^26,42,44,62^, evidenced by the quantitative match between subjects’ accuracy and RTs with model fits (data points vs. solid gray lines in Fig. 4b-c; R^2^, 0.97±0.01 for accuracy and 0.98±0.01 for RTs, mean±s.e.m across subjects). After estimating the model parameters, we used them to predict the shape of the psychophysical kernel for the 0% coherence motion energies used in the experiment. These predicted kernels (Fig. 4e-f, solid gray lines) closely matched the experimentally derived ones (R^2^, 0.57), establishing that the dynamics of the kernels were both qualitatively and quantitatively compatible with stationary sensory weights and a decision-making process based on bounded accumulation of evidence.

In a second experiment, we focused on a more complex sensory decision that required combining multiple spatial features over time (Fig. 5a). Subjects categorized faces based on their similarity to two prototypes. Each face was designed to have only three informative features (eye, nose, and mouth) (Fig. 5b). On each trial, the mean strengths (percent morph) of these three features were similar and randomly chosen from a fixed set spanning the morph line between the two prototypes. However, the three features fluctuated independently along their respective morph lines every 106.7 ms (Fig. 5c; see Methods). All other parts of the faces remained fixed halfway between the two prototypes during and across all trials and, therefore, were uninformative. Further, each frame of the face stimulus was quickly masked to prevent subjects from consciously perceiving the small fluctuations in eyes, nose, and mouth. Subjects reported the identity of the face (closer to prototype 1 or 2) with a saccadic eye movement to one of the two targets, as soon as they were ready. Therefore, we could measure both the choice and RT. The key difference with the direction discrimination task was that instead of one stimulus attribute that fluctuated over time (motion energy), there were three attributes that fluctuated independently. The three informative features could support the same or different choices in each stimulus frame and across frames. This task provided a richer setting to test how humans combine multiple spatial features to make a decision.

**Figure 5.**
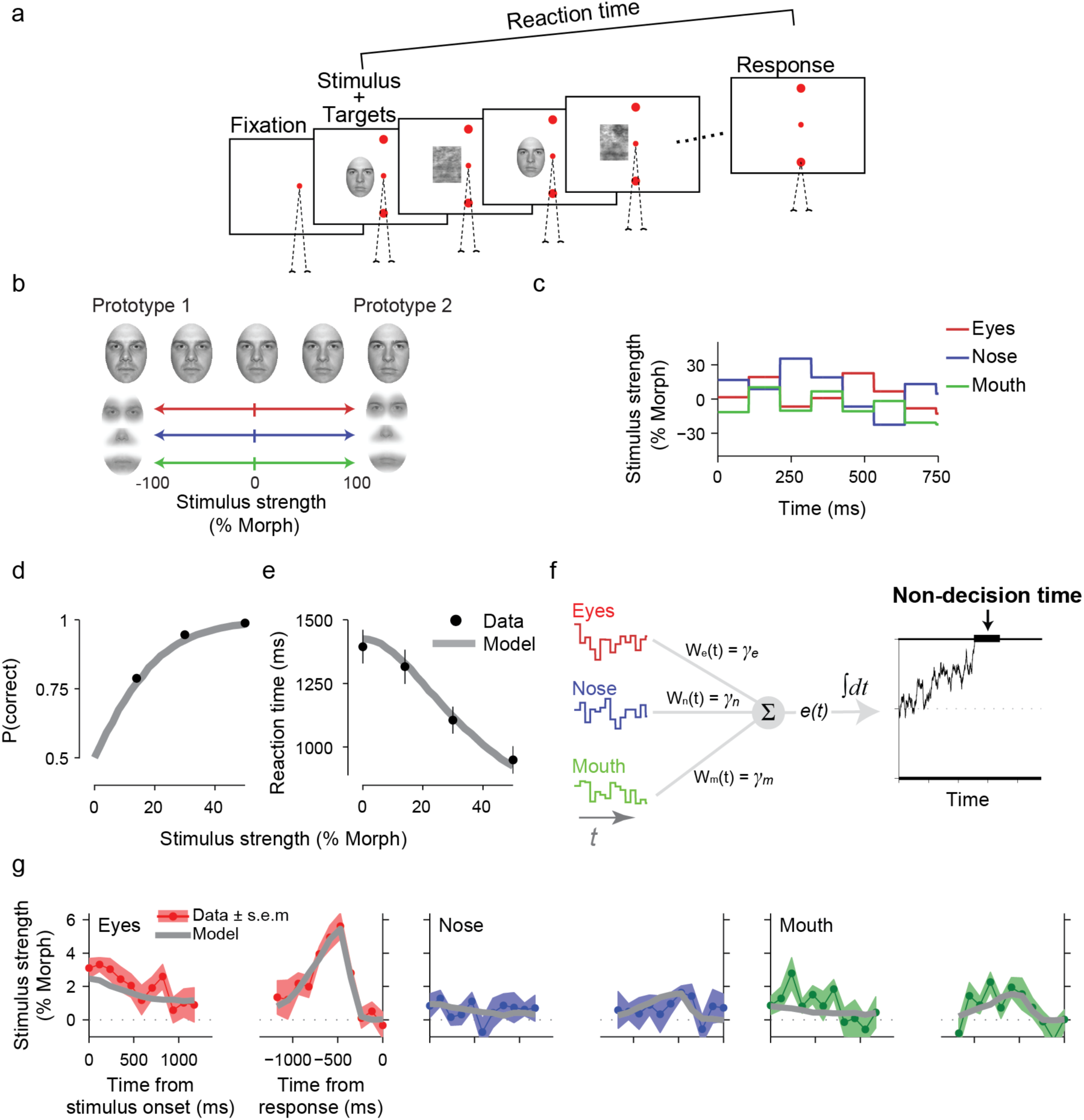
Psychophysical reverse correlation in a face discrimination task with multiple informative features reveals relative weighting of features and kernel dynamics similar to the direction discrimination task. (**a**) Task design. Subjects viewed a sequence of faces interleaved with masks and reported whether the face identity matched one of two prototypes. They reported their choice with a saccadic eye movement to one of the two targets, as soon as ready. (**b**) Using a custom algorithm, we designed intermediate morph images between the two prototype faces such that only three facial features (eyes, nose, and mouth) could be informative. These features were morphed independently from one prototype (+100% morph) to another (−100% morph), enabling us to create stimuli in which different features could be biased toward different identities. All regions outside the three informative features were set to half-way between the prototypes and were uninformative. (**c**) The three informative features underwent subliminal fluctuations within each trial (updated with 106.7ms interval). The mean morph levels of the three features were similar but varied across trials. Fluctuations of the three features were independent (Gaussian distribution with standard deviations set to 20% morph level). (**d-e**) Choice accuracy increased and RTs decreased with stimulus strength. Data points are averages across subjects. Error bars are s.e.m across subjects. Gray lines are model fits. (**f**) The DDM model used to fit subjects’ choices and RTs extends the model in Fig. 2 by assuming different sensitivity for the three informative features. Momentary evidence is a weighted average of three features where the weights correspond to the sensitivity parameters. The momentary evidence is integrated toward a decision bound. (**g**) Psychophysical kernels estimated from the model (gray lines) match subjects’ kernels for the three features. Shaded areas are s.e.m across subjects.

Consistent with the simpler direction discrimination task, as the average morph level of the three features increased and the stimulus better resembled one of the prototypes, choices became both more accurate and faster (Fig. 5d-e). The psychophysical kernels of the three features (Fig. 5g) had rich dynamics. First, the eye kernels had larger amplitude than the mouth and nose kernels, suggesting that choices were more strongly influenced by fluctuations in the eye region^68,69^. Second, the stimulus-aligned kernels dropped gradually over time, and the saccade-aligned kernels showed a characteristic peak a few hundreds of milliseconds prior to the choice. Had we not introduced these characteristic dynamics earlier, one might have been tempted to interpret the kernel dynamics as changes in spatiotemporal weighting of facial features for our task. However, as we explained above, such kernel dynamics could also arise from a decision-making process based on fixed weights and bounded accumulation of evidence.

Therefore, we explored whether a multi-feature integration process with stationary weights for eyes, nose, and mouth regions could quantitatively explain our results. For each stimulus frame, the model calculated a weighted sum of the three features to estimate the momentary sensory evidence and then integrated this momentary evidence over time in a bounded diffusion model (Fig. 5f, see Methods). Fitting the model to the choice and RT distributions provided a quantitative match for both (gray lines in Fig. 5d-e are model fits; R^2^, 0.998±0.001 for accuracy and 0.98±0.01 for RTs) and the resulting parameters led to kernels that well matched the dynamics of experimentally observed kernels for the three features (R^2^, 0.74).

Overall, bounded integration of sensory evidence during decision-making introduces characteristic dynamics in the psychophysical kernels that quantitatively match the data both for simple, one-dimensional sensory decisions (direction discrimination), and for more complex, multi-dimensional decisions (face discrimination). Knowing these signature dynamics enables experimenters to understand their results in a more comprehensive framework that accommodates nuances of both sensory and decision-making mechanisms. More generally, a model that is fit to choices and RTs can generate exact predictions about the shape and time course of psychophysical kernels. Comparison of these predictions against experimentally derived psychophysical kernels, as we did above, provides a powerful test for the validity of models, beyond those offered by psychometric and chronometric functions.

## Testing for temporal dynamics of sensory weights

Our exploration of the model and fits to experimental data in the previous sections focused largely on cases in which sensory weights were static and the dynamics of the psychophysical kernel were solely due to the decision-making process. However, as discussed earlier, changes of sensory weights could also be a major factor in shaping psychophysical kernels (e.g., Figs. 3c, S1, and S2). In theory, a model-based approach to understanding kernel dynamics should be able to distinguish changes of sensory weights from decision-making processes because of their distinct effects on the choice and RT distributions. To test this prediction, we simulated a direction discrimination experiment in which decisions were made by accumulation of weighted sensory evidence toward a bound in the presence of non-decision time and various dynamics of sensory weights (Fig. S5). Then, we used the simulated choice and RT distributions to fit an extended DDM that allowed temporal dynamics of sensory weights. The model recovered the weight dynamics and accurately predicted psychophysical kernels of the simulated experiments in each case (Fig. S5). A few thousand trials, similar to those available in our experimental datasets, were adequate to achieve accurate fits and predictions. Therefore, there does not seem to be critical limitations in the ability of a model-based approach to detect sensory weight dynamics, when such dynamics are present.

Knowing about the model’s ability, we extended the DDMs used in the previous section to explore dynamics of sensory weights for human subjects that participated in the direction discrimination and face discrimination tasks. The extended models included linear and quadratic terms to capture a wide variety of temporal dynamics (Eq. 23 and Eq. 25 for direction discrimination and face discrimination, respectively). The results did not support substantial temporal dynamics of sensory weights in either task (12 out of 13 subjects of the direction discrimination task and all subjects of the face discrimination task showed static weights). Overall, the addition of temporal dynamics to the weight function did not significantly improve the fits or the match between model and experimental psychophysical kernels (for direction discrimination, Eq. 23, *β*_1_, −3.0±1.6, p=0.10, median, −0.65, and *β*_2_, −2.1±2.3, p=0.36, median, 0.17; for face discrimination, Eq. 25, *β*_1_, −0.19±0.10, p=0.09, median, 0.10, and *β*_2_, 0.055±0.028, p=0.08, median, 0.028). These results about static sensory weights are in agreement with previous electrophysiological recordings in area MT, which suggest largely stable neural responses during stimulus viewing^70^. Because similar models could accurately recover weight dynamics in the simulated data, we do not think our observation about the experimental data is caused by a low power for the detection of weight dynamics in the model or a fundamental bias to attribute changes of psychophysical kernels to the decision-making process.

## Characteristic dynamics of psychophysical kernels for decision bound, noise, input correlation, inhibition, and leak in the decision-making process

Although a simple DDM for accumulation of evidence captures several key aspects of behavior in sensory decisions, it is only an abstraction for the more complex computations implemented by the decision-making circuitry. More complex and nuanced models are required both to explain details of behavior and to create biologically-plausible models of integration in a network of neurons. We use this section to explore a non-exhaustive list of key parameters commonly used in various implementations of evidence integration models. For clarity, all simulations are for models without non-decision time to isolate the effects of these model parameters from the effects of non-decision time.

First, we focus on how changes of decision bound influence the shape of psychophysical kernels. The effect is best demonstrated by our mathematical proof that the kernel is proportional to sensory weights in a simple DDM without non-decision time (Eq. 2). The constant of this proportionality is 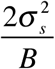, where *B* is the bound height and 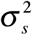 is the variance of stimulus fluctuation. As a result, if subjects increase the decision bound to improve their accuracy^23,47,71^, psychophysical kernels will shrink (Fig. S6a-b). Conversely, when they reduce the decision bound to emphasize speed over accuracy, the kernels are amplified. These changes are expected because a lower decision bound boosts the effect of stimulus fluctuations on choice and vice versa. This scaling with bound height can be corrected by estimating the decision bound from behavior and multiplying the kernels by it, as we did for the experimental results in the previous sections.

Changes of decision bound, however, are not limited to adjustments of speed-accuracy tradeoff across trials. They can also happen within a trial. Recordings from the frontoparietal neurons that represent integration of evidence supports the presence of a stimulus-independent signal that pushes the integration process towards the decision bound^23,24,42,44^. This urgency signal can be formulated as a reduction of decision bound over time (Fig. 6a)^44,72^. Therefore, a strong urgency signal would lead to an inflation of the psychophysical kernel, as shown in Fig. 6b. The intuition behind kernel inflation with urgency is similar to that explained above. A growing urgency gives small stimulus fluctuations a higher chance to cause a bound-crossing event, increasing the influence of later stimuli on the choice. This larger influence shows itself as a rise in the psychophysical kernel.

**Figure 6.**
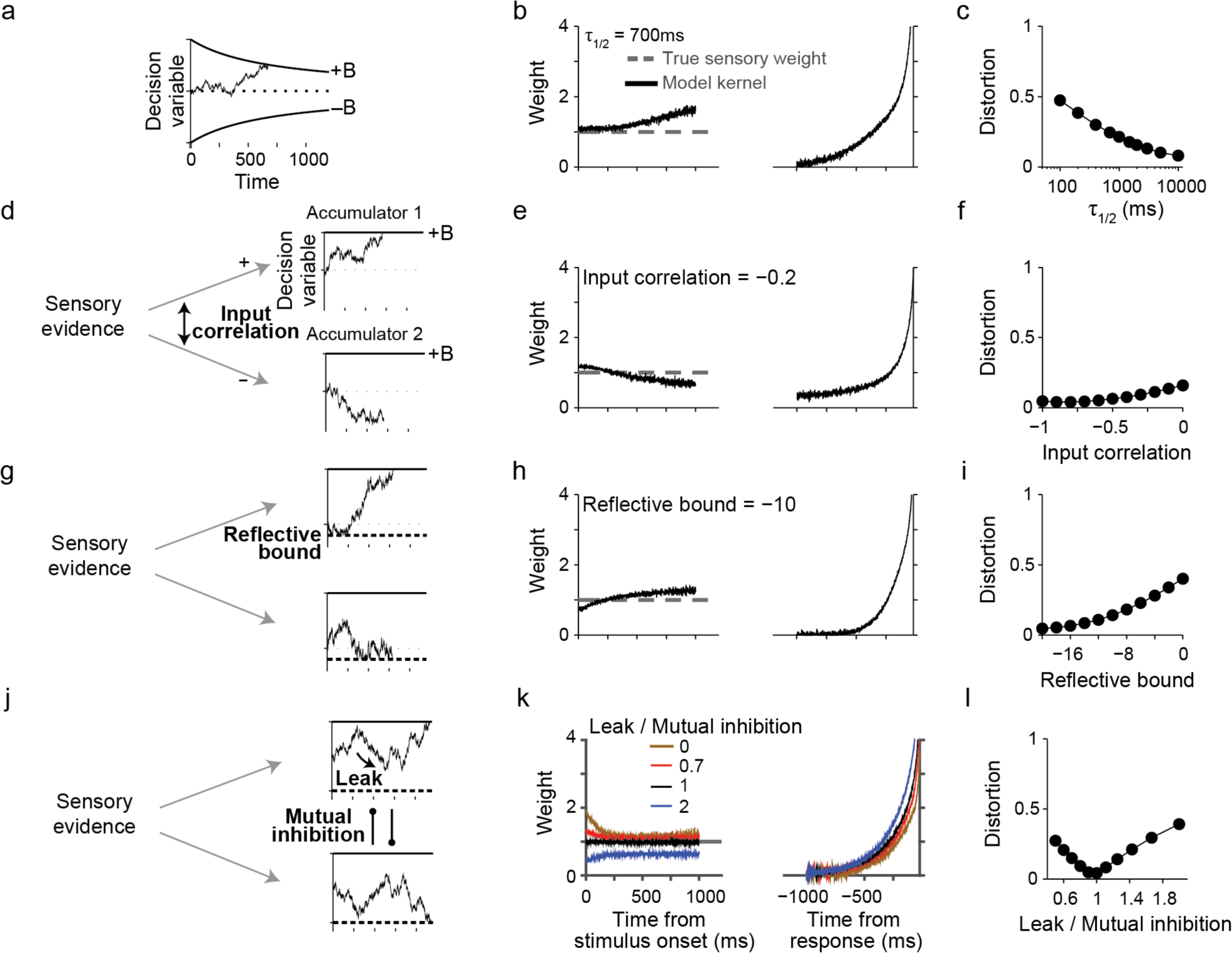
Psychophysical kernels are susceptible to changes of decision bound, input correlation, mutual inhibition, integration time constant, and limited dynamic range. The figure shows extensions of DDM and systematic deviations that additional realism to the model can cause in psychophysical kernels. Conventions are similar to Fig. 3, except that we focus only on RT tasks. Also, to isolate the effects of different model parameters from the effect of non-decision time, we use zero non-decision time in these simulations. (**a-c**) Collapsing decision bound (urgency signal) inflates psychophysical kernel over time. The rate of bound collapse is defined by *τ*_1/2_ — the time it takes to have a 50% drop in bound height. (**d-f**) Extending DDM to a competition between two bounded accumulators reveals that input correlation of the accumulators has only modest effects on psychophysical kernels, causing an initial overshoot followed by an undershoot compared to true sensory weights. (**g-i**) The presence of a lower reflective bound in the accumulators causes an opposite distortion: an initial undershoot followed by a later overshoot. (**j-l**) Balancing the effect of mutual inhibition by making the integrators leaky causes the model to behave like a DDM, eliminating the effects of both the inhibition and leak on the psychophysical kernels (black curves in m). Any imbalance between leak and inhibition, however, causes systematic deviations in the kernels from the true sensory weights (brown, red, and blue curves in k). See Fig. S7 for more examples.

The proportionality constant explained above 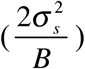 also points at another interesting possibility in which changes of stimulus variance over time, if present, could systematically distort psychophysical kernels: larger stimulus noise inflates the kernel (Fig. S6c-e). It is customary in most experiments to keep stimulus noise fixed. However, one should be aware of the possibility that fluctuations in the stimulus noise across trials^73^ or a time-varying stimulus noise within trials may distort behaviorally measured psychophysical kernels, as suggested by our equations. This contrasts with the effects of internal (neural) noise for the representation of sensory stimuli or the DV. We show in Methods that in a bounded DDM, internal noise does not have a systematic effect on psychophysical kernels of RT tasks (but compare to unbounded DDM in Methods).

Electrophysiological recordings from motor planning regions of the primate brain suggest that integration of sensory evidence is best explained with an array of accumulators, rather than a single integration process^44,49,50,74–80^. A class of models that match this observation better than the simple DDM is competing integrators—one for each choice—that accumulate evidence toward a bound^20,25,26,50,79,81–83^. Our mathematical proof for the shape of the kernel does not exactly apply to these models. However, many of these models can be formulated as extensions of the DDM with new parameters added to provide more flexible dynamics^22^. For example, a DDM is mathematically equivalent to two integrators that receive perfectly anti-correlated inputs (correlation = −1) and, consequently, are anti-correlated with each other^22,26^. We use these links and our understanding of the psychophysical kernels in the DDM to provide intuitions for kernel distortions in more complex models with competing integrators.

One can adjust the noise correlation in the input of the two competing integrators to achieve a more realistic neural implementation of the evidence integration. Neurons representing different choices tend to be negatively correlated^84^ but it is rare for them to be perfectly anti-correlated. Perfect anti-correlation in neural responses is not expected because even when signal correlations are negative, noise correlations tend to be close to zero or slightly positive^85–87^. Fig. 6d-f show that the shape of the psychophysical kernel is only minimally affected by a wide range of correlations in the input of two competing integrators. Sizeable distortions arise only when the input correlation approaches 0, in which case the kernel is initially inflated but later drops below the true sensory weight (Fig. 6e and S7a). The inflation is caused by an effective increase in the diffusion noise because for low input correlations the noise in either integrator can facilitate a bound crossing. However, as time passes, the mean DV of unterminated decisions becomes increasingly more negative due to diffusion noise and attrition of trials whose DV exceeds the bound. The more negative DVs reduce the effect of new input for determining the outcome of the process, shrinking the kernel below the true sensory weights.

Another commonly used feature of a biologically plausible implementation of the integration process is a lower reflective bound that limits how low the DV of each integrator can go^21,25–27,50^. Such reflective bounds are inspired by the observation that the spike count of neurons is limited from below and cannot become negative. When the reflective bounds are far enough from the starting point of the integrators they tend to have only a modest effect on the psychophysical kernel (Fig. 6g-i and S7b). However, their effect grows quickly as the reflective bound approaches the starting point. Unlike the input correlation, reflective bounds cause the psychophysical kernel to begin lower than the true sensory weight but exceed it later. The initial underestimation happens because reflective bounds limit movements below the starting point and, thus, reduce both the effective noise and the effective counter-evidence for each choice. Later in the integration process, the integrators are on average closer to their decision bounds compared to a model without lower reflective bounds. This amplifies the psychophysical kernel because input fluctuations are more likely to lead to a decision bound crossing.

Another parameter to consider is mutual inhibition between the integrators. Several models incorporate such inhibition either through direct interactions between the integrators^25,50^ or indirectly through intermediate inhibitory units^21,53,88^. When the activity of one integrator grows, it suppresses the other one, creating winner-take-all dynamics that amplify the difference of the two integrators and effectively prevents the losing integrator from gaining the upper hand^21,53,88^. This suppression can cause dramatic distortions in psychophysical kernels, especially early in the integration process, because the mutual inhibition magnifies the effect of early sensory evidence on the state of the two integrators (Fig. 6j-l brown lines and S7c). As the integration process continues and the losing integrator drops far enough from its decision bound to exert significant inhibition on the other integrator, the behavior of the model converges to a simple DDM and the kernel converges on the true sensory weights.

Mutual inhibition is often combined with decay (leak) in the integration process (Fig. 6j) to create richer dynamics and curtail the effects of inhibition^22,25^. The balance between the leak and mutual inhibition in the model defines whether it implements bistable point attractor dynamics or line attractor dynamics^22^. When mutual inhibition dominates (leak/inhibition ratio<1), psychophysical kernels show an early amplification but later converge on the true sensory weights (Fig. 6k, red lines), for the same reasons explained in the previous paragraph. When leak and inhibition balance each other out, the model acts similar to a line attractor and the psychophysical kernels resemble those of a DDM (Fig. 6k, black lines). Finally, when leak dominates, the integrators lose information and decisions are influenced less by input fluctuations, especially for early sensory evidence in the trial. Consequently, stimulus-aligned psychophysical kernels systematically underestimate the sensory weights (Fig. 6k, blue lines). However, the dynamics of the kernel qualitatively resemble the true sensory weight, except for the earliest times. On the other hand, the response-aligned kernels are distorted and accelerated compared to a DDM, reflecting the shorter integration time constant and stronger influence of later evidence on the decision^25,32^.

The presence of bias in the decision-making process is another factor that can cause distortions in psychophysical kernels. Two competing hypotheses have been suggested for implementation of bias in the accumulation to bound models. One hypothesis is a static change in the starting point of the accumulation process (or an equivalent static change in decision bounds)^40,47,48,89^, which would cause an initial inflation in the psychophysical kernels without a lasting effect (Fig. S8a). A second hypothesis is a dynamic bias signal that pushes the decision variable toward one of the decision bounds and away from the other^45^. This dynamic bias signal can be approximated by a change in the drift rate of DDM, which would cause a DC offset in the psychophysical kernels (Fig. S8c).

Overall, when the parameters of the decision-making process are adjusted to implement linear integration of sensory evidence, non-decision time and changes of decision bound (urgency) have the largest effects on the kernel. Input correlation, lower reflective bounds, and balance of leak and inhibition tend to have smaller effects, unless these parameters take extreme values. Interestingly, and perhaps by luck, applying these more complex models to the experimental data of the previous section resulted in model parameters that closely resembled linear integration of evidence, which is why the DDMs in the last section performed so well. Because of this parameterization, these more sophisticated models would produce predictions similar to the simple DDM about the dynamics of the psychophysical kernel. However, we note that this observation may not generalize to other experiments and should, therefore, be tested for new behavioral paradigms on a case-by-case basis.

More generally, different parameters of decision-making models have different and even opposing effects on the expected shape of psychophysical kernels. As a result, a mixture of these features can, in principle, generate a variety of kernel dynamics, depending on their exact parameters. To illustrate this point, we consider models with two competing integrators that have different levels of mutual inhibition, leak, collapsing bounds, and sensory and motor delays (Fig. 7a). For static sensory weights over time, this class of models can generate monotonically decreasing kernels (Fig. 7b), monotonically increasing kernels (Fig. 7c), or kernels that exactly match the true sensory weights (Fig. 7d), depending on the model parameterization. To understand this diversity, consider for example the opposing effects of collapsing bounds (urgency) and non-decision time on the kernels. The gradual reduction of the kernel due to non-decision time can cancel out the increase of the kernel due to urgency. Alternatively, one of the two effects may overpower the other one. Complementary to the examples in Fig. 7, one can also imagine parameterizations that would result in a flat psychophysical kernel in the presence of non-stationary true sensory weights. The presence of mutual inhibition and leak further complicate the relationship of sensory weights and psychophysical kernels and expand the space of possible dynamics for the kernels.

**Figure 7.**
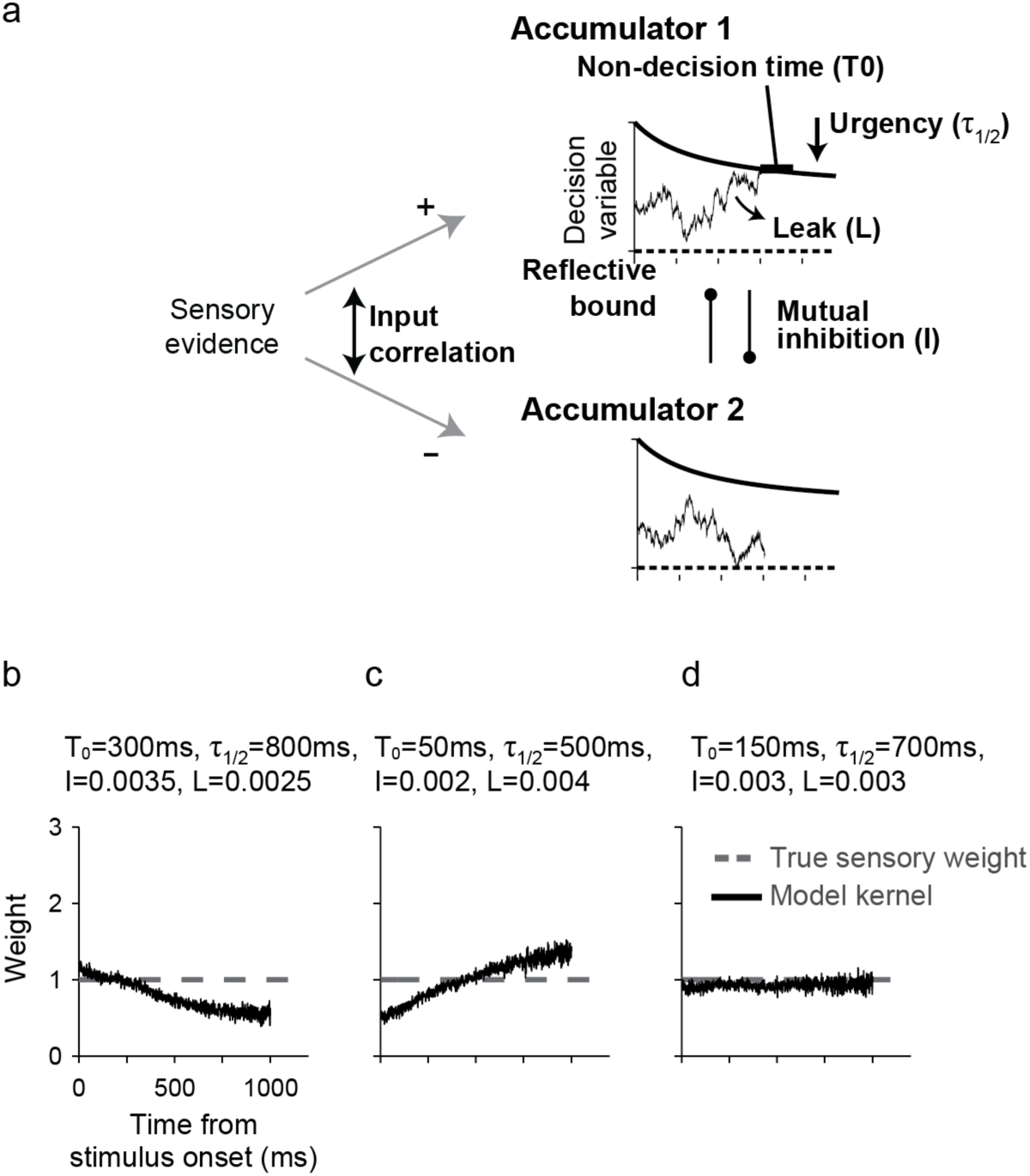
A decision-making model that has a mixture of parameters with opposing effects on psychophysical kernels can create a diversity of kernel dynamics for static sensory weights. (**a**) A model composed of two competing integrators that allows different ratios of leak and inhibition, collapsing decision bounds, and nondecision times. The model also has input correlation >−1 and reflective lower bounds, but they are fixed for simplicity. (**b**) When bound collapse is small and non-decision times are long, the kernel drops monotonically over time. (**c**) When bound collapse is large and non-decision times are short, the kernel rises monotonically. (**d**) When these opposing factors balance each other, the kernel becomes flat.

This diversity of plausible kernel dynamics without meaningful fluctuations in true sensory weights cautions about ignoring the mechanisms of the decision-making process for interpretation of experimentally derived psychophysical kernels. One needs a model-based approach to decipher the true meaning of fluctuations in psychophysical kernels and the information they impart about spatiotemporal filters of sensory processes.

## Discussion

A key goal of systems neuroscience is to explain behavior as a sequence of neural computations that transform sensory inputs to appropriate motor outputs. For perceptual decisions, this sequence includes sensory processes that form neural representations of a stimulus in sensory cortices and decision-making processes that plan the best choice based on these sensory representations. Psychophysical reverse correlation has been originally developed to infer sensory filters that approximate sensory processes^3–5^. Recent studies, however, have begun to use the technique for inferring the properties of the decision-making process^31,32,37,64,90,91^. Here, we show that an isolated perspective is vulnerable. To ensure correct interpretation of psychophysical kernels, one has to adopt an integrative perspective that sees psychophysical kernels as a product of both sensory and decision-making processes.

We arrived at our conclusion through a systematic exploration of how psychophysical reverse correlation is influenced by the decision-making process and under what conditions it provides a good approximation of sensory filters. We showed that neither the integration of sensory evidence nor the termination of the integration process by reaching a decision bound fundamentally limits the recovery of sensory weights. However, nuances that are fixtures of real experiments but often receive little attention can be major sources of deviation between psychophysical kernels and sensory weights. Examples include sensory and motor delays or input correlation of the integration process, which cause kernels with a downward trend unrelated to sensory weights, or urgency and lower reflective bounds in the integration process, which can cause upward trends in the kernels. Previous theoretical explanations of reverse correlation have ignored these nuances and have caused confusion about what can be gleaned from psychophysical kernels. We also showed that the kernels are susceptible to how the integration process is implemented. Bistable point attractor or line attractor dynamics, which are implemented through different combinations of mutual inhibition, self-excitation, and decay of activity (leak) in neural networks, yield different kernel dynamics. We conclude that psychophysical kernels are influenced by both sensory and decision-making processes and show how they can be used to provide information about both types of processes.

Making the interpretation of psychophysical reverse correlation dependent on the decision-making process is likely to face opposition because of the historical influence of signal detection theory^8,9^ and the fact that under the assumptions of SDT, reverse correlation matches the true sensory weights. However, we note that this match is misleading. In particular, SDT explains the dynamics of psychophysical kernels by simplifying the decision-making process and shifting its complexity to the sensory processes. This shift is inaccurate both because it depends on unsubstantiated sensory processes and because it ignores the known complexity in the decision-making process. The perspective offered by SDT is also insufficient because it fails to explain changes of psychophysical kernels that stem from flexibility of the decision-making process. For example, setting the speed and accuracy of choices^23,24^ or adjustment of behavior following feedback^42,92,93^ often depends on rapid alterations in the decision-making process. These alterations also change psychophysical kernels, as explained in Results. The integrative framework proposed here would correctly identify the source of changes, whereas SDT would have to misattribute them to changes of sensory weights.

Proper partitioning of the contribution of sensory and decision-making processes enables testing hypotheses about neural mechanisms of behavior. We provide key signatures of psychophysical kernels for a variety of different mechanisms and implementations. Matching these signatures to the patterns observed in an experiment enables hypothesis formation. To quantitatively test those hypotheses, one can implement computational models of sensory and decision-making processes, fit them to some aspect of behavior, and then generate predictions about another aspect of behavior. For example, by fitting the DDM to the distribution of choices and RTs, we could accurately predict the dynamics of psychophysical kernels in our experiments. Further, our mathematical proofs and simulations provide a comprehensive framework for predicting changes of psychophysical kernels under various experimental manipulations. For example, when a manipulation leads to improved accuracy, one can use our method to distinguish the two potential sources of the improved performance: increased sensitivity (e.g., attentional mechanisms) or changes of decision bound (speed-accuracy tradeoff). Whereas increased sensitivity is expected to increase the magnitude of the psychophysical kernel, increased decision bound or reduced urgency is expected to reduce the kernel magnitude. Such contrasting predictions highlight the ability of psychophysical reverse correlation to separate models that may not be easily distinguished by more conventional measurements, including changes in psychometric function or its derivatives such as overall accuracy.

However, psychophysical kernels on their own are inadequate to determine the nuances of the sensory and decision-making processes. As we show in Figs. 6 and 7, multiple mechanisms can have similar effects on the shape of psychophysical kernels. Some of these mechanisms with opposing effects on the kernels can even cancel each other out and lead to a flat kernel, complicating a model-free interpretation of experimentally derived kernels. To reduce interpretational errors, one should always assess psychophysical kernels in conjunction with the choice and RT distributions. Mechanisms that have a similar effect on the kernel often have contrasting effects on choices and RTs. For example, variability of non-decision times, input correlation, and mutual inhibition can all lead to stimulus-aligned kernels that shrink with time. However, these three mechanisms (i) have different effects on the shape of the tail of RT distribution, (ii) make different predictions about accuracy for the same sensitivity function (Fig. S9a), and (iii) differ in the exact shape of the psychophysical kernels (note the small but detectable, qualitative and quantitative differences in the dynamics of the kernels in Fig. S9a). The combination of these signatures enables separation of different mechanisms. A similar contrast in kernel dynamics and RT and choice distributions exists for dropping bounds and reflective bounds, which both cause an inflation of psychophysical kernels over time. Along the same lines, when combinations of different mechanisms with opposing effects lead to a flat kernel (e.g., Fig. 7d), the distribution of RT and choice is often different compared to when the kernel is flat because none of those mechanisms contribute to the decision-making process or a different combination of parameters flattens the kernel (Fig. S9b). A three-pronged approach based on the shape of the kernel, and RT and choice distributions makes a powerful technique for uncovering the mechanisms that shape behavior. However, there are also important limitations on what and how much can be learned just from behavior. For example, when tails of RT distribution play a key role in distinguishing different mechanisms, the amount of available data and presence or absence of experimental manipulations that may encourage fast responses will influence our ability to arrive at the correct conclusions.

To maximize the information gained from psychophysical kernels, it is important to set up the model fitting and model prediction in such a way that minimizes overlap between the fitted and predicted aspects of the data. For example, using stimulus fluctuations on individual trials to predict the choice and calculating model kernels for the same stimulus fluctuations is unlikely to provide new insights because any model that fits the choices well could also replicate the kernels. However, by leaving the specific stimulus fluctuations aside for fitting the choices or by predicting kernels for a non-overlapping group of trials one can reveal potential discrepancies between the model and data.

A key piece of information for proper interpretation of psychophysical kernels is to know which part of the stimulus is used for the decision-making process. Fixed duration tasks with long stimulus durations are generally unsuitable because the start and termination times of the decision-making process are opaque to the experimenter. When subjects commit to a choice by integration of sensory evidence toward a bound, they tend to ignore later evidence, causing a downward trend in the kernels. However, if the integration process also begins at variable times, the downward trend can be masked. Overall, the presence or absence of temporal dynamics in psychophysical kernels in fixed-duration tasks does not have a unique interpretation. It is more suitable to use tasks in which stimulus duration is controlled by the experimenter and varies randomly from trial to trial because they enable the experimenter to determine which part of the stimulus is used for the decision. However, one should be careful in using those designs because just the variability of stimulus duration in itself can introduce temporal dynamics in kernels (Eq. 14). Reaction time task designs are most suitable because they minimize ambiguity about which part of the stimulus was used for the decision-making process.

We also note that tasks that use very brief stimulus presentations are not immune to the influence of the decision-making process on psychophysical kernels. Brief stimulus presentations are often used to infer the spatial structure of sensory filters at the cost of ignoring their temporal dynamics. However, brief stimulus presentations do not guarantee instantaneous decisions. Several studies have demonstrated that accumulation of evidence to a bound is at work even for brief stimuli^94,95^, as evidenced by large RT differences for different stimulus strengths. Even brief stimulus presentations produce extended trails of activity in sensory and action-planning neurons^60,96,97^. An extended decision-making process for a briefly presented stimulus makes the kernel susceptible to the deviations explained above. For example, a change in speed-accuracy tradeoff can amplify the inferred spatial filters without a real change in sensory processes.

In light of our results, past studies that relied on psychophysical reverse correlation can teach us more than most of them were designed for. It is still valid to interpret psychophysical kernels as the best approximation of a linear-nonlinear model, such as SDT, to the behavior. However, it is important to keep in mind that such an interpretation is about effective associations between sensory information and choice^98^, which should not be confused with the spatiotemporal filters that shape sensory representations or readout of sensory representations to form the evidence used in the decision-making process. It is also important to note that these effective associations do change under various conditions that do not change sensory representations (e.g., changes of decision bound), while the model-based approach suggested here is likely to recover the true dynamics of sensory weights. The results of our paper do not refute careful use of psychophysical reverse correlation in past studies. Rather, we try to elevate psychophysical reverse correlation from a technique that reveals only effective associations of stimulus and choice to a technique that reveals the inner working of the sensory and decision-making processes that underlie the choice.

Using a fixed-duration design, several studies have found monotonically decreasing psychophysical kernels^32,36,54,99^. Kernel dynamics in those studies could have a sensory origin or be due to static sensory weights and a decision-making mechanism that terminates according to some criterion (e.g., Fig. 3g). It has been common to assume one possibility and ignore the other. Testing these two possibilities explicitly is likely to yield new insights and trends across experiments. Several other studies have reported a flat psychophysical kernel in fixed-duration designs and interpreted it as a signature of perfect integration of sensory evidence. This interpretation could be correct if subjects set their decision bound too high to reach during stimulus viewing, as shown in Brunton et al.^33^. However, a static kernel could also arise from a variety of sensory and decision-making mechanisms and does not uniquely support perfect integration of sensory evidence. Also, in addition to the mechanisms explained in Results, a static kernel in a non-RT task could be produced with probabilistic sampling of evidence rather than its integration. In general, it is best not to rely solely on qualitative patterns of psychophysical kernels. As we suggest above, these qualitative signatures should be just a starting point for building mechanistic hypotheses, which should then be tested with detailed, quantitative modeling of behavior and electrophysiological studies of its underlying neural responses.

## Methods

We examine how well psychophysical reverse correlation recovers sensory weights in perceptual tasks where decisions are based on accumulation of sensory information^25,48,100,101^. First we prove that the core computations for integration of evidence or termination of the decision-making process based a decision bound do not cause any deviation of psychophysical kernels from true sensory weights. Then, we demonstrate that non-decision times, urgency, reflective bounds, and interactions between accumulators are substantial sources of deviation. Finally, we show how one can accurately interpret the complexity of psychophysical kernels by explicit modeling of both sensory and decision-making processes.

### Psychophysical reverse correlation for bounded accumulation models

Drift diffusion models are commonly used to approximate integration of evidence in two-alternative sensory decisions^17–19^. In these models, a weighting function is applied to momentary sensory evidence and the result is integrated over time until the integrated evidence (the decision variable, DV) reaches either an upper (positive) or a lower (negative) bound. Each bound corresponds to one of the choices. The sensory weighting function is assumed constant in some experiments such as direction discrimination of random dots where sensory neurons show more or less constant activity proportional to stimulus strength throughout the stimulus presentation period^70^. However, the exact form of the weighting function is usually unknown in most experiments and could change dynamically depending on context. We prove below that in the absence of sensory and motor delays, reverse correlation accurately recovers the weights applied to sensory evidence in a drift diffusion model (DDM).

In a reaction time task, where subjects report their choices as soon as ready, the psychophysical kernel is

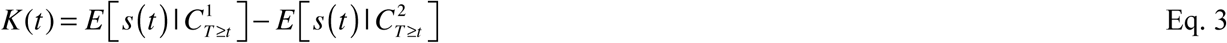

where 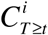 indicates all trials in which choice *i* is made at times equal or larger than *t*, and 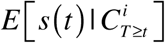 is the average stimulus at time *t* conditional on the choice being made at a later time (*T* ≥ *t*). The intuition for this formulation is that a trial contributes to the calculation of the kernel at time *t* only if the choice on that trial is not recorded by the experimenter before *t*. For an unbiased decision-maker and a stimulus distribution symmetric around zero, 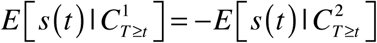, leading to 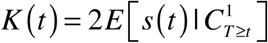. Therefore, we need to calculate only one of the two conditional averages.

In a DDM, the choice is made through integration of sensory evidence:

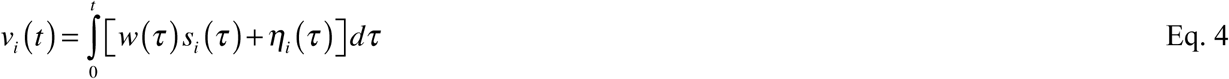

where *v_i_*(*t*) indicates the DV on trial *i* at time *t*, *w*(t) is the weight applied on the stimulus at times *τ* ≤ *t*, *s_i_*(*τ*) are stimuli sampled from a Gaussian distribution with mean 0 and variance 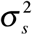, and *η_i_*(*τ*) represents internal (neural) noise for the representation of sensory and integration processes. We assume that the internal noise does not bias the representation and can be approximated with a Gaussian distribution with mean 0 and variance 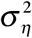.

Because integration continues until *v_i_*(*t*) reaches one of the two bounds (+*B* or −*B*),

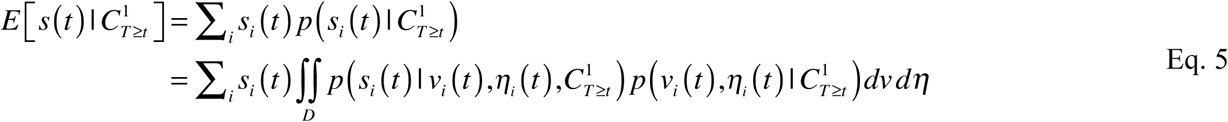

where the integration domain *D* is [−*B*,+*B*]. Using Bayes rule and by plugging Eq. 5 in Eq. 3, we get

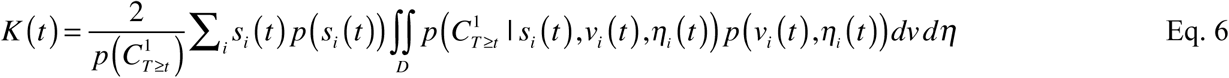

where 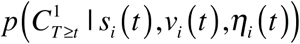 is the probability of reaching the upper bound after time *t*, when the decision maker observes stimulus *s_i_*(*t*) and has an existing decision variable *v_i_*(*t*). This bound-crossing probability has an analytical solution in DDM^48^. Note that after observing stimulus *s_i_*(*t*), the distance of the accumulated evidence from the upper bound is 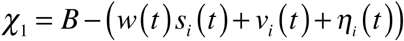, and the distance from the lower bound is 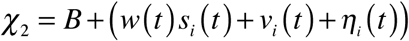. Because *s*(*t*) and *η*(*t*) have zero mean, the overall drift is zero and the bound-crossing probability is:

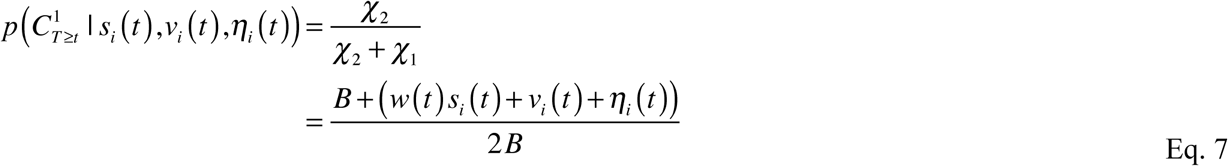

Therefor, Eq. 6 can be written as:

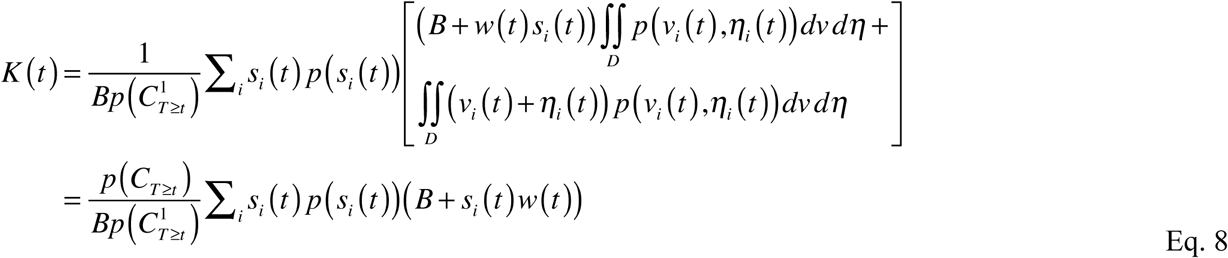

where *p*(*C_T_*_≥_*_t_*) is the combined probability of reaching either of the two decision bounds after time *t*. The second equality in Eq. 8 stems from two equations. First,

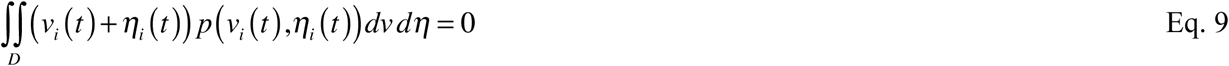

because for a neutral stimulus, the DV is symmetrically distributed around the starting point. Second,

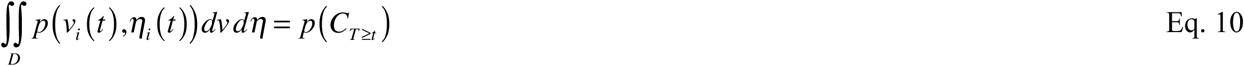

because 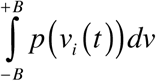 reflects the total probability of the DV between the decision bounds at *t*, and because this unabsorbed probability mass is guaranteed to be fully absorbed by the decision bounds in finite time *T* ≥ *t*^102^.

For an unbiased decision-making process and a stimulus with zero mean 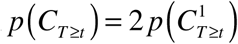. As a result, Eq. 8 simplifies to

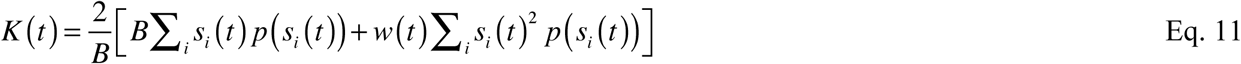

For a Gaussian stimulus with zero mean, 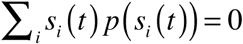 and 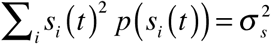. Therefore,

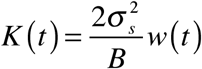

This equation, which we highlight in Results (Eq. 2), shows that the result of psychophysical reverse correlation is proportional to the sensory weights. The proportionality constant is 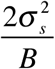, which explains how reverse correlation is modulated by properties of the stimulus (stimulus variance) and parameters of the decision-making process (decision bound). Eq. 2 also shows that psychophysical kernels are independent of internal noise in a DDM. Internal noise does not cause a systematic deviation in estimated kernels, although it could affect the confidence interval of the estimated kernels in real experiments, where a limited number of trials are available for measuring the kernels.

Based on Eq. 2, the outcome of psychophysical reverse correlation is expected to change if the decision bound is not constant. For example, urgency in the decision-making process is often equivalent to a drop in the decision bound^42,44,72^, which should lead to a gradual increase of the reverse correlation kernel even in the absence of changes in sensory weights (Fig. 6a-c). Note that we define urgency as an additive signal for competing accumulation processes, which under certain conditions (e.g., anti-correlated input to the accumulators) can be translated to collapsing bounds in the DDM. This is different from the alternative definition of urgency based on gain in the accumulation process^43,61^, which cannot be easily translated to a bound change in DDM.

The proof above holds only if the choice can be recorded as soon as the decision-making process terminates. In practice, one has to take into account sensory and motor delays that postpone initiation of action. These delays imply that the stimuli presented immediately before the behavioral response do not influence the choice^52,62^. As a result, the psychophysical kernel drops to zero prior to the choice. Due to trial-to-trial variability of these delays, it is not possible to know purely based on behavior, which part of the stimulus did or did not contribute on any individual trial, but on average one can expect a descending trend in psychophysical kernel close to the time of the response (Fig. 3i-m).

In addition to the bound height and non-decision time, other factors can cause deviation of psychophysical kernels from sensory weights. DDM is a simplified model of the more complex computations implemented by the neural circuit that underlies the choice. In the simplest case, one should consider an array of accumulators that interact and compete with each other^21,31,50,88,103,104^, forcing us to consider correlation between accumulators, mutual inhibition, and leak which can cause systematic deviations in the kernels (Fig. 6d-f, 6j-l). Further, real neurons do not accommodate negative firing rates. A lower reflective bound in each accumulator can introduce additional systematic biases in the kernel (Fig. 6g-i). A closed form, mathematical solution for the psychophysical kernel in the presence of all these factors is complex and beyond the scope of this paper. Therefore, we use simulations to explore the parameter space of different model variations and demonstrate how different factors change the kernel (see “Model simulation” below).

### Psychophysical reverse correlation for unbounded accumulation of evidence in fixed-duration tasks

If decisions in a fixed-duration task are made by unbounded integration of evidence psychophysical kernels will correctly reflect the dynamics of sensory weights (Fig. 3a-c, S1). The proof is as follows. If integration begins with stimulus onset and continues for the whole stimulus duration, *T_s_*, the decision variable at the end of the stimulus in trial *i* will be

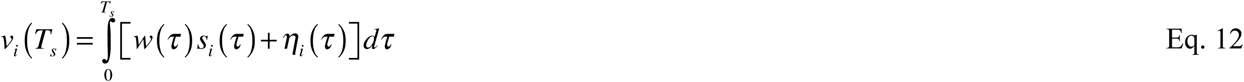

If the sensory input, *s*(*t*), is drawn from a Gaussian distribution with mean 0 and variance 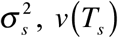 will have a mean of 0 and variance 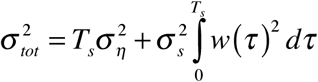. The model selects choice 1 for the positive DV and choice 2 for the negative DV. Therefore, the psychophysical kernel will be

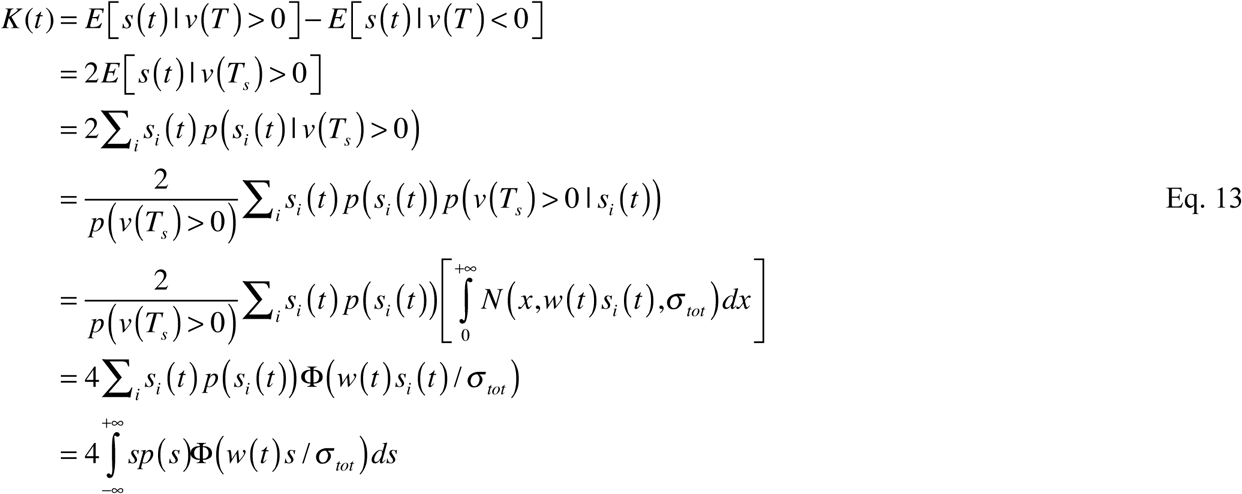

where Φ(*x*) is the cumulative distribution function of a standard normal probability density function with mean 0 and standard deviation 1. The last equality in the equation is due to the i.i.d. property of *s*(*t*) within and across trials.

The kernel equation can be further simplified as

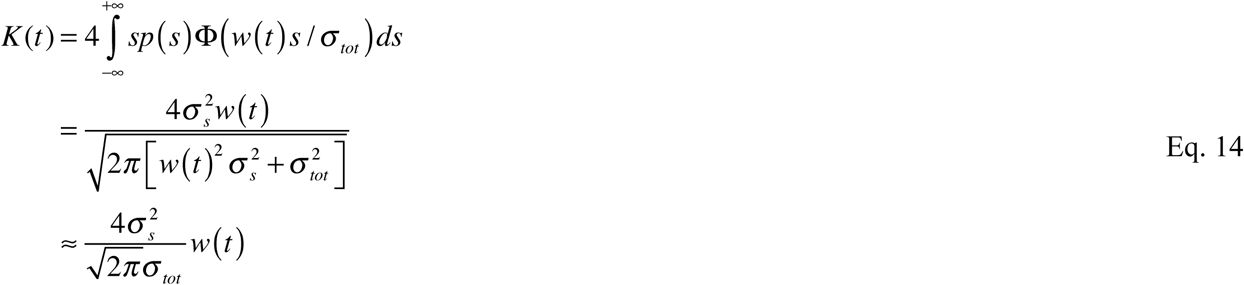

where the approximation in the last line is based on 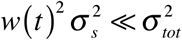, which is usually true unless stimulus durations are very short.

Based on Eq. 14, the kernel is proportional to the sensory weight function, and the constant of proportionality scales with stimulus variance 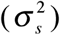, similar to Eq. 2 for bounded accumulation.

However, note that the kernel is inversely proportional to *σ_tot_*, which is a function of the stimulus duration. Dependence of kernels on stimulus duration calls for caution in interpretation of results when a mixture of stimulus durations are used in an experiment. The kernel for each stimulus duration is scaled differently, inducing artificial dynamics in the average kernel across all durations.

Even when stimulus durations are the same across trials, subjects can begin integration at variable times across trials and commit to a choice at different times during stimulus viewing, causing temporal dynamics in the kernel that do not reflect the true dynamics of sensory weights (see Results and Discussion for a more detailed explanation).

## Model simulation

We simulated four different classes of bounded accumulation models: (i) DDM without non-decision time, (ii) DDM with non-decision time^18,28,48^, (iii) DDM with non-decision time and urgency^24,42,44^; and (iv) Competing accumulators with different input correlation, reflective bound, mutual inhibition, and leak^20,25,26,50,79,81,82^.

For each class and parameter combination, 10^6^ trials were simulated to obtain an accurate estimation of the psychophysical kernel. Sensory input for each trial was a sequence of independent draws from a Gaussian distribution with mean 0 and *σ_s_* = 1. This input was multiplied with a weight function, *w*(*t*), which could be constant (Fig. 3, 6, 7, S3, S6–9) or vary over time (Fig. S1-2, S5). This weight function dictated the significance that each sample played in shaping the decision. The outcome (termed momentary evidence) was passed to each integration model to calculate the DV and generate a choice for each trial. As explained above, the presence or absence of internal noise did not bear strongly on the measured kernels in RT tasks, as long as enough trials were available for the measurement. We calculated psychophysical kernels based on the simulated choices and sensory stimuli according to Eq. 3 and compared the results against the weight function used in the simulation (Fig. 3, 6, 7, S1-2, S6-9).

*Drift diffusion models*. In these models momentary evidence was integrated over time until the DV reached a lower bound (−*B*) or an upper bound (+*B*), which corresponded to the two choices. For models without non-decision time, integration stopped immediately and a choice was registered after the bound crossing. For models with non-decision time, the integration process stopped after bound crossing but the choice was registered after a random time, drawn from a Gaussian distribution. The stimuli presented between bound crossing and the choice were included in the calculation of psychophysical kernels to emulate realistic experimental conditions, where experimenters do not know the exact non-decision time on each trial. The mean and standard deviation of the distribution of non-decision time in Fig. 3j-l and S2 were 300 ms and 100 ms, respectively, compatible with past studies^28,41,42,61^. In Fig. 3m, the mean varied between 0 and 800ms, and the standard deviation was equal to 1/3 of the mean. In Fig. S3, we tested the effects of different means, variance, and skewness of the non-decision time distribution on the measured psychophysical kernels.

For DDMs with urgency, we reduced the decision bound according to a hyperbolic function:

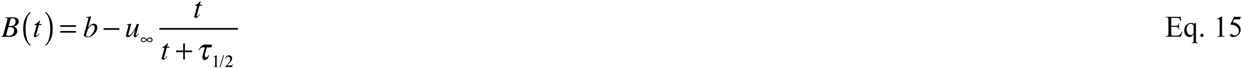

where *b* is the initial bound height, *u*_∞_ is the asymptotic reduction in bound height, and *τ*_1/2_ is the time to reach 50% of the reduction. We set *b* = 60, *u*_∞_ = 60, and *τ*_1/2_ = 400*ms* for Fig. 6b. In Fig. 6c, *τ*_1/2_ varied while the other model parameters were kept constant.

For the simulation of fixed-duration tasks (Fig. 3b, 3g, 3l, S1), we incorporated past experimental observations that the decision-making process could effectively stop before the termination of the stimulus^32,51,54,91^. Stimulus durations were 1 s on all trials. The full stimulus duration was used for the calculation of psychophysical kernel to reflect the standard practice and experimenters’ lack of knowledge about the exact time of the decision on each trial.

*Competing accumulator models*. DDM is a low-parameter model and by design lacks the sophistication of a biologically plausible neural network that implements the integration process^21,105–107^. A more biologically plausibility alternative is a model with a bank of accumulators that interact and compete with each other^50,108^. For a two-alternative decision, the simplest instantiation of such a model has two accumulators, each integrating sensory evidence in favor of one of the two choices^25,26,88^. The model reaches a decision when one of the accumulators crosses its bound. In addition to the DDM parameters (bound height and non-decision time), the competing accumulator model has the following parameters (see Eq. 16):

1. Input correlation (*ρ*) determines the correlation between sensory inputs of the two accumulators. The inputs are explained by a two-dimensional Gaussian distribution with mean 0 and covariance matrix 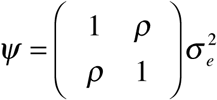, where 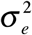 reflects the combined variance of weighted stimulus noise and internal noise.
2. The second parameter is a reflective bound (*R*) that defines a lower limit for the DV of each accumulator.
3. The third parameter is the strength of inhibitory interactions between the accumulators (*I*). This mutual inhibition is widely assumed to be a key component of biological circuits of decision-making and a key factor in shaping neuronal response dynamics^21,25,104^. When *I* > 0, the strength of mutual inhibition for accumulators 1 and 2 at time *t* is *Iv_2_ (t*) and *Iv_1_* (*t*), respectively, where *v*_1_ and *v*_2_ are the DVs of the two accumulators. Because the magnitude of inhibition is proportional to the accumulated evidence, even small *I* can have dramatic effects on the decision-making process.
4. The fourth model parameter is “leakage” in the integration process (*L*). In the absence of mutual inhibition, the leak makes the model behave as an Ornstein-Uhlenbeck process^102^, causing the DVs to decay faster as they get farther from their starting point. In the presence of mutual inhibition, the balance of leak and inhibition creates a variety of attractor dynamics. When the leak and inhibition parameters are equal (*L* = *I*), the difference of the DVs of the two accumulators implements a DDM: a line attractor that reflects the accumulated difference of momentary evidence of the two accumulators^22^. When mutual inhibition exceeds leak (*L* < *I*), a saddle point emerges in the state space of the model, which exponentially amplifies small initial differences of the DVs of the two accumulators over time. This amplification boosts the effect of early stimulus fluctuations on the decision^25^. Conversely, when the leak parameter exceeds mutual inhibition (*L* > *I*), a point attractor emerges in the state space, causing differences in the DVs of the two accumulators to decay over time. This decay reduces the effect of early stimulus fluctuations on the choice.

The equation that governs our simulations of competing accumulator models in Fig. 6, 7, and S7 is:

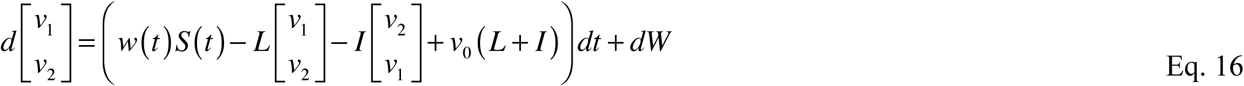

where *dv* denotes the change in *v* over a small time interval *dt*, *L* is the leak term, *I* is the mutual inhibition, and *v*_0_ is the starting point of decision variables. *S*(*t*) is a vector that represents the sensory inputs to the two accumulators. For the simulations in Fig. 6, 7, and S7, we assumed that that two accumulators were driven in opposite directions by the input stimuli, that is 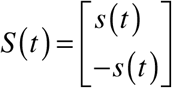 is a 2D Gaussian noise term with mean 0 and covariance *ξdt*, where 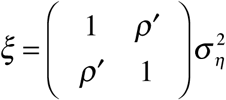. We adjusted *ρ*′ to achieve a desired input correlation (*ρ*) as defined above. *v*_1_ and *v*_2_ started at *v*_0_ · *v*_0_ (*L* + *I*) created a stable point at *v*_0_. The decision variables were subjected to two non-linearities: a lower reflective bound (*R*) and an upper absorbing bound (*B*).

Fig. 6e-f demonstrate distortions in the psychophysical kernels for different input correlations (*ρ* was set to −0.2 in Fig. 6e and varied between −1 and 0 in Fig. 6f; *B* = 30, *R* = −∞, *I* = 0, *L* = 0, 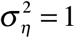, *v*_0_ = 0). In the absence of a lower reflective bound, inhibition or leak, the model became mathematically equivalent to a DDM whenever *ρ* = −1. In Fig. 6h-i, we tested the effect of a lower reflective bound (*R* was set to −10 in Fig. 6h and varied between −20 and 0 in Fig. 6i; *ρ* = −1, *B* = 30, *I* = 0, *L* = 0, 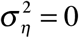, *v*_0_ = 0). Fig. 6k-l show how the balance of leak and mutual inhibition distorted psychophysical kernels. For these simulations, we kept *L* + *I* = 0.006 and systematically changed the ratio 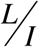 between 0.5 and 2 (*ρ* = −1, *B* = 60, *R* = 0, = 0, *v*_0_ = 30). We also show the shape of the kernel in the absence of a leak (brown lines, *I* was set to 0.003). For these simulations the lower reflective bound was set to 0 to ensure that negative DVs in one accumulator did not excite the other accumulator. To best isolate the effect of individual parameters of the model, we set the non-decision time and urgency to zero in Fig. 6d-l. Fig. 7 shows the effect that conjunctions of different parameters have on the psychophysical kernel. The standard deviation of non-decision time was set to 1/3 of its mean in this figure.

*Comparison of model kernels and sensory weights*. For each model we calculated the psychophysical kernel as explained by Eq. 3. To directly compare the kernels with the sensory weights implemented in the model, we divided the kernels by the scaling factor of Eq. 2 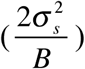. For models with dynamic bounds (Fig. 6a-c, 7), we used the average bound height from the stimulus onset to the median RT to calculate the scaling factor. For unbounded models (Fig. 3a-c), we used the scaling factor in Eq. 14 (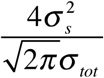). After scaling the kernels and making them comparable to the sensory weights, we quantified the difference between the stimulus-aligned weight and kernel functions using root mean square error:

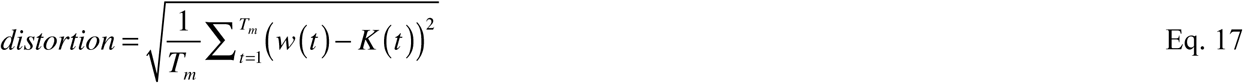

where *T_m_* is the stimulus duration in simulations of fixed duration tasks or the median RT in simulations of RT tasks.

### Psychophysical tests

We performed two experiments to test our model predictions: direction discrimination with random dots, and a novel face discrimination task. All subjects were naïve to the purpose of the experiments and provided informed written consent before participation. All procedures were approved by the Institutional Review Board at New York University. Throughout the experiments, subjects were seated in an adjustable chair in a semi-dark room with chin and forehead supported before a CRT display monitor (refresh rate 75Hz, viewing distance 52-57 cm). Stimulus presentation was controlled with Psychophysics Toolbox^109^ and Matlab. Eye movements were monitored using a high-speed infrared camera (Eyelink, SR-Research, Ontario). Gaze positions were recorded at 1kHz.

*Direction discrimination*. Thirteen human subjects performed a RT version of the direction discrimination task with random dots^26,28,42^. Data from six subjects have been previously reported in Kiani et al.^26^ and data from the remaining subjects have been reported in Purcell & Kiani^42^. Both studies used a similar trial structure. Subjects initiated each trial by fixating a small red point at the center of the screen (FP, 0.3° diameter). After a variable delay, two targets appeared on the screen, indicating the two possible motion directions. Following another random delay, the dynamic random dots stimulus^110^ appeared within a 5-7° circular aperture centered on the FP. The stimulus consisted of three independent sets of moving dots shown in consecutive frames. Each set of dots was shown for one video frame and then replotted three video frames later (Δ*t* = 40 ms; density, 16.7 dots/deg^2^/s). When replotted, a subset of dots were offset from their original location (speed, 5 deg/s), while the remaining dots were placed randomly within the aperture. The percentage of coherently displaced dots determined the strength of motion. On each trial, motion strength was randomly chosen from one of six possible values: 0%, 3.2%, 6.4%, 12.8%, 25.6%, 51.2% coherence. Subjects reported their perceived direction of motion with a saccadic eye movement to the choice target in the direction of motion. Once the motion stimulus appeared, subjects were free to indicate their choice at any time. RT was recorded as the difference between the time of motion onset and eye movement initiation. For the calculation of psychophysical kernels, motion energy of the random dot stimulus was calculated for each trial over time (see below).

*Face discrimination task*. Nine human subjects performed a novel experiment designed to test our model predictions for more complex decisions on multi-dimensional sensory stimuli. Subjects reported the identity of a face on each trial as soon as they were ready (Fig. 5a). The stimuli were drawn from morph continuums between photographs of two faces (MacBrain Face Stimulus Set^111^, http://www.macbrain.org/resources.htm). We developed a custom algorithm that morphed different facial features (regions of the stimulus) independently between two prototype faces. Our algorithm started with 97 manually-defined anchor points on each face and morphed one face into another by linear interpolation of the positions of anchor points and textures inside the tessellated triangles defined by the anchor points. The result was a perceptually seamless transformation of the geometry and internal features from one face to another. The anchor points also enabled us to morph different regions of the faces independently. We focused on three key features (eyes, nose, and mouth) and created independent series of morphs for them. The faces that were used in the task were composed of different morph levels of these three features. Anything outside those features was set to the halfway morph between the prototypes. The informativeness of the three features (stimulus strength) was defined based on the mixture of prototypes, spanning from −100% when the feature was identical to prototype 1 to +100% when it was identical to prototype 2 (Fig. 5b). 0% morph corresponded to the middle of the morph line, where the feature was equally shaped by the two prototypes.

By varying the three features independently we could study spatial integration through creating ambiguous stimuli in which different features could support different choices. We could also study temporal integration of features by varying the three discriminating features within each trial (Fig. 5c). The three discriminating features for each stimulus frame were drawn from independent Gaussian distributions. The mean and standard deviation of these distributions were equal and fixed within each trial, but the means varied randomly from trial to trial. We tested seven mean stimulus strengths (−50%, −30%, −14%, 0%, +14%, +30%, and +50% morph level). The standard deviation was 20% morph level. Sampled values that fell outside the range [−100% +100%] (0.18% of samples) were replaced with new samples inside the range.

Changes of the stimulus within a trial were implemented in a subliminal fashion such that subjects could not consciously perceive variation of facial features and yet their choices were influenced by these variations. We achieved this goal using a sequence of stimuli and masks within each trial. The stimuli were morphed faces with a particular combination of the three discriminating features. The masks were created by phase randomization of the intermediate face between the two prototypes^112^. For the majority of subjects (7/9), each stimulus was shown without a mask for one monitor frame (13.3ms). Then, it gradually faded out over the next 7 frames as a mask stimulus faded in. For these frames the mask and the stimulus were linearly combined, pixel-by-pixel, according to a half-cosine function, such that in the last frame the weight of the mask was 1 and the weight of the stimulus was 0. Immediately afterwards, a new stimulus frame with a new combination of informative features was shown, followed by another cycle of masking, and so on. For a minority of subjects (2/9), we replaced the half cosine function for the transition of stimulus and mask with a full cosine function, where each 8-frame cycle started with a mask, transitioned to an unmasked stimulus in frame 5, and transitioned back to a full mask by the beginning of the next cycle. We did not observe any noticeable difference in the results of the two presentation methods and pooled their data. The masks ensured that subjects did not perceive minor changes in key features over time within a trial. In debriefings following each experiment, all subjects noted that they saw one face in each trial but the face was covered with various masks over time.

## Analysis of behavioral data

*Motion energy*. Due to the stochastic nature of the random dot motion stimuli, the strength of motion of a stimulus with a fixed coherence fluctuated from one frame to another. We quantified these stimulus fluctuations by calculating motion energy^66^. Details are described elsewhere^32^. Briefly, we used two pairs of spatiotemporal filters, each selective for one of the two motion directions discriminated by the subject. Each direction-selective filter was formed by summation of two space-time separable filters. The spatial filters were even and odd symmetric fourth-order Cauchy functions:

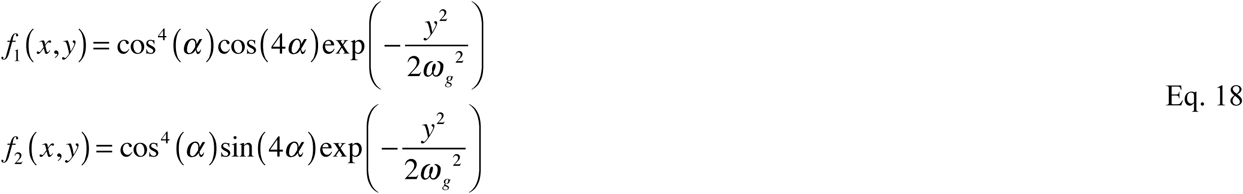

where 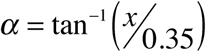 and *ω_g_* = 0.05. The two temporal filters were:

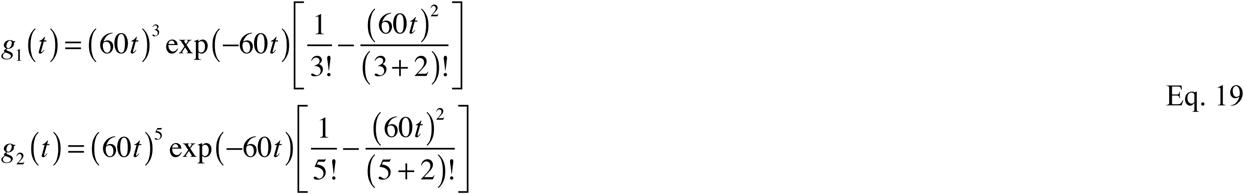

The two pairs of direction-selective filter were constructed by combining the two spatial filters with the two temporal filters: *f*_1_*g*_1_ + *f*_2_*g*_2_ and *f*_2_*g*_1_ − *f*_1_*g*_2_ were selective for one motion direction, whereas *f*_1_*g*_1_ − *f*_2_*g*_2_ and *f*_2_*g*_1_ + *f*_1_*g*_2_ were selective for the opposite direction. The parameters of Eq. 18 and Eq. 19 were chosen to (i) match spatial and temporal bandpass properties of MT neurons, (ii) to maximize selectivity of the directional filters for the speed of coherent motion in the stimulus (5 deg/s), and (iii) to reproduce the width of direction-selectivity tuning curves of MT neurons. We convolved the 3D spatiotemporal pattern of the stimulus in each trial with these four filters, squared the results, and then summed them for each pair of filters to measure local motion energies at each stimulus sub-region over time. The local energies were summated across space and subtracted from the energy of the opposing pair of filters to obtain fluctuations of the net motion energy in one direction over time.

Average motion energies increased linearly with stimulus coherence (Fig. S4b). However, the lag in the temporal filters caused the effect of stimulus fluctuations to show up in the motion energies with ~50ms delay (Fig. 4d and S4a), as shown before^32,66^.

*Psychophysical kernels*. For the direction discrimination task, we used motion energies of 0% coherence trials to perform reverse correlation (Eq. 3) on the responses of human subjects (3389 trials). For Fig. 4e-f, we first computed each subject’s psychophysical kernel and then averaged the kernels across subjects. Each subject’s stimulus- and response-aligned kernels were calculated up to the subject’s median RT, ensuring that at least half of trials contributed to the calculations. When averaged across subjects, the kernels were shown up to the shortest median RT. For the response-aligned kernels (Fig. 4f), we rounded the RT to the onset of the last stimulus frame on the monitor. The temporal resolution of kernels (13.3ms) was dictated by the refresh rate of the monitor (75 Hz).

For the face discrimination task, we used fluctuations of eyes, nose, and mouth morph levels in the 0% morph trials to calculate psychophysical kernels of individual features (Fig. 5c) (3530 trials). Similar conventions to the direction discrimination task were used for averaging kernels across subjects and plotting them, except that because of the longer stimulus frame durations in the face discrimination task (106.7ms), the kernels were temporally coarser.

We did not perform any smoothing of the psychophysical kernels of the two tasks to avoid obscuring their dynamics.

## Model fits to the behavioral data and prediction of psychophysical kernels

*Direction discrimination task*. We used a simple DDM to fit subjects’ choices and RTs in order to predict their psychophysical kernels. The model had four degrees of freedom: decision bound height (*B*), mean non-decision time (*T*_0_), standard deviation of non-decision time 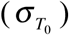, and a sensitivity parameter (*γ*). The sensitivity parameter determined the mean of momentary evidence (*μ* = *γC*) conferred by a motion stimulus with coherence *C*. The bound height and sensitivity were in units of the standard deviation of the momentary evidence per unit time (*σ_e_*), which we set to 1. This formulation of DDM, which has been used widely in the past, directly maps to the formulation presented earlier in methods:

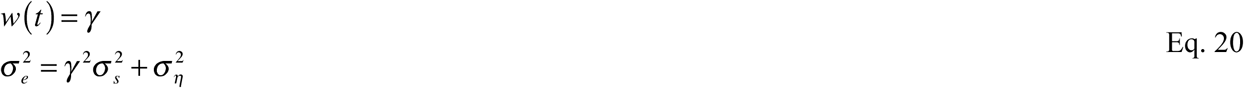

The probability of crossing the upper and lower decision bounds at each decision time was calculated by solving the Fokker-Planck equation^102,113^:

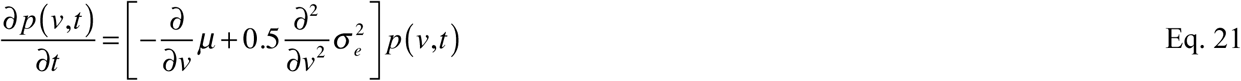

where *p*(*v, t*) is the probability density of the DVs at different times. The boundary conditions were

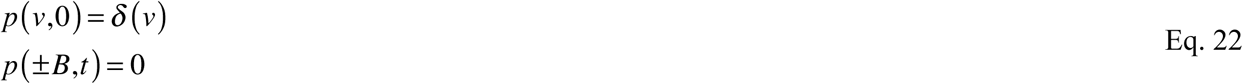

where *δ*(*v*) denotes a delta function. The first condition enforced that the DV always started at 0 and the second condition guaranteed that the accumulation terminated when the DV reached one of the bounds. RT distributions for each choice were obtained by convolving the distribution of bound crossing times with the distribution of non-decision times (Fig. 2).

We fit model parameters by maximizing the likelihood of the joint distribution of the observed choices and RTs in the experiment. For a set of parameters, the model predicted a distribution of RTs for each possible choice for the stimulus strength used in each trial. These distributions were used to calculate the log-likelihood of the observed choice and RT on single trials. These log-likelihoods were summed across trials to search for the best set of model parameters that maximized this sum. The model parameters were fit separately for each subject. To avoid local maxima, we repeated the fits from 10 random initial points and chose the parameters that maximized the likelihood function. Fig. 4b-c show the average fits across subjects. The high quality of fits for individual subjects and the average subject indicated that the DDM provided an adequate explanation for the computations underlying behavior in the direction-discrimination task. Compatible with past studies, adding urgency to the model, or replacing the DDM with a competing accumulator model did not fundamentally change the fits because the parameterization of these more complex models stayed in a regime that approximated the line attractor dynamics of the DDM^22,26^.

To test if a time-varying weighting function provided a better fit to the behavioral results, we modified Eq. 20 by adding linear and quadratic temporal modulations to the drift rate:

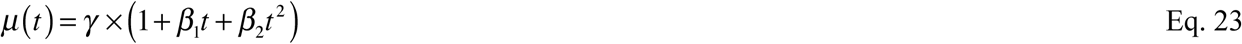

where *β*_1_ and *β*_2_ are additional degrees of freedom in the model.

*Prediction of psychophysical kernels in the direction discrimination task*. We used the best model parameters that fit the RT and choice distributions to predict subjects’ psychophysical kernels. We use the term prediction because moment-to-moment fluctuations of motion energies were not used to fit the model parameters and the fitting procedure did not create any explicit link between these fluctuations in the stimuli used in the experiments and single trial choices and RTs. Using the model parameters we predicted choices and RTs for 10^5^ simulated trials with 0% motion coherence and calculated motion-energy kernels for the model choices. Because the sensitivity parameter of the model was calculated for motion coherence, we first divided motion energies by the slope of the line that related average motion coherence to the average motion energy (Fig. S4). This division converted motion energy fluctuations within a trial to equivalent stimulus coherence fluctuations, which were directly passed to the model to generate a choice and a reaction time for each simulated trial. We used these choices and RTs to calculate the model prediction for the kernels and superimposed them on subjects’ kernels (Fig. 4e-f).

*Face discrimination task*. We extended the simple DDM explained above to include three different sensitivity parameters for the three facial features (*γ_e_* for eyes, *γ_n_* for nose, and *γ_m_* for mouth), increasing the total number of parameters to 6 (the other parameters were *B*, *T*_0_, and 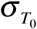). The mean momentary evidence at each time in a trial was:

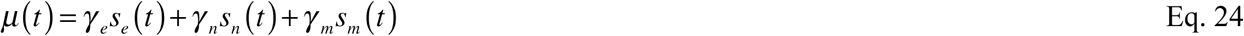

where *s_e_*(*t*), *s_n_*(*t*), and *s_m_*(*t*) were the morph levels of eyes, nose, and mouth at time *t* on the trial. Note that *μ*(*t*) is a time-varying drift rate based on the exact fluctuations of stimulus strengths on individual trials, unlike the drift rate in the model for the direction discrimination task. Our goal was to obtain the relative sensitivity for the three informative facial features. Because the average morph levels of the three features were identical in each trial, using the average morph to derive the drift rate would have made the three sensitivity parameters redundant. The fitting procedure to subjects’ choices and RTs were as explained for the direction discrimination task, except that we used Eq. 24 to include a time-varying drift rate, *μ*(*t*). Also, note that because we used the exact stimulus fluctuations in the Fokker-Plank equation, *σ_s_* was excluded from the definition of noise 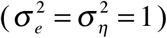. The model parameters were fit separately for each subject using the maximum-likelihood procedure explained above.

To test if a time-varying weighting function provided a better fit to the behavioral results, we modified Eq. 24 to allow linear and quadratic temporal modulations to the drift rate, similar to what we did for Eq. 23:

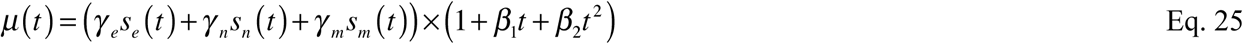

*Model psychophysical kernels for the face discrimination task*. The procedure for deriving the model kernels was similar to that for the direction discrimination task. We simulated 10^5^ trials with 0% morph and passed the fluctuations of the three informative features to get model choices and RTs for individual trials. We then used these choices to calculate the model kernels for the three features and superimposed the result on subjects’ psychophysical kernels for comparison (Fig. 5g). Because we used the stimulus fluctuations for fitting the model parameters, the kernels derived from the model were not pure predictions, unlike the direction-discrimination task. However, note that the model kernels were not directly fit to match the data either. They were calculated based on an independent set of simulated 0% morph trials, making the comparison in Fig. 5g informative.

## Acknowledgement

We thank Michael Waskom and John Sakon for insightful comments on the manuscript. This work was supported by the National Institutes of Health Grant R01-MH109180, a Sloan Research fellowship, and a McKnight Scholar Award to RK. LS was supported by NRSA training grant R90-1R90DA043849, BP was supported by a post-doctoral fellowship from the Simons Collaboration on the Global Brain, and GO was supported by post-doctoral fellowships from Japan Society for the Promotion of Science and from Charles H. Revson Foundation.

## Supplementary Figures

**Figure S1.**
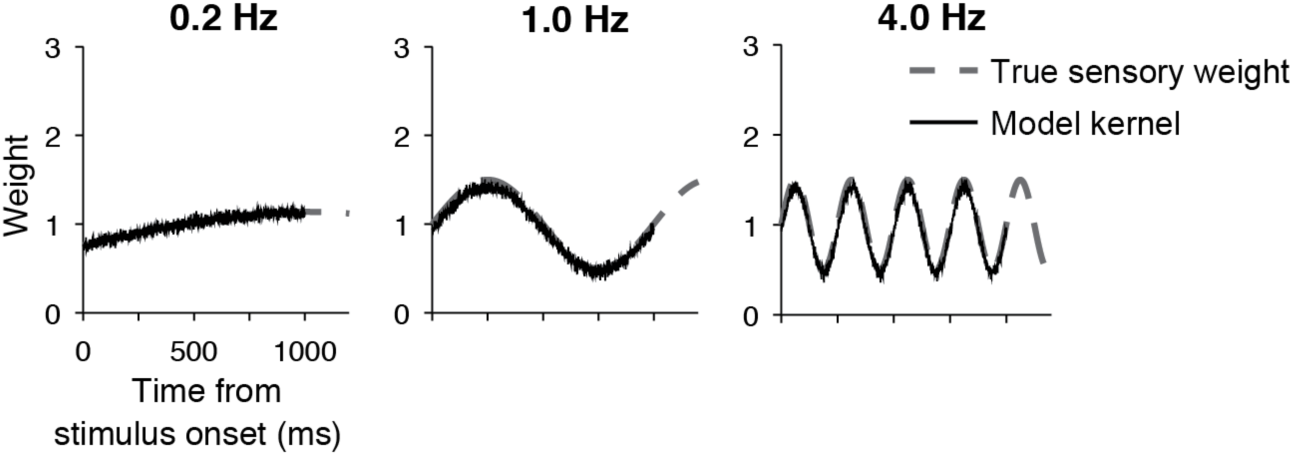
For an unbounded DDM, psychophysical kernels recover the true sensory weights. Related to Fig. 3a-b. This figure shows three simulated models with sensory weights that fluctuated at 0.2, 1, and 4 Hz. Any frequencies of weights could be accurately recovered by the psychophysical kernel. See Fig. 3c for the quantification of kernel distortions as a function of temporal frequency of weights.

**Figure S2.**
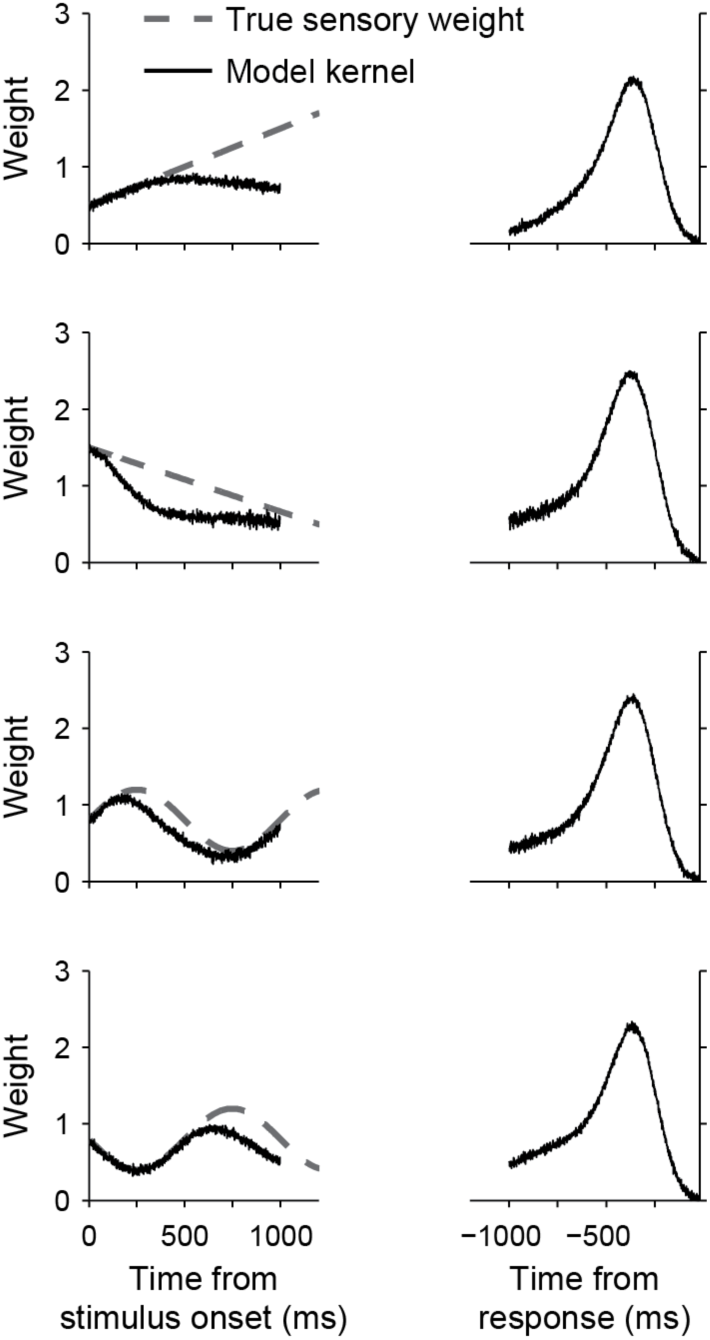
Psychophysical kernels deviate from sensory weights because of incomplete knowledge about decision time. Related to Fig. 3i-m. The figure shows four simulated bounded DDMs with non-decision time and time varying sensory weights. The mean and standard deviation of non-decision time are set to 300ms and 100ms, respectively, and decision bound is set to 30. The non-decision time causes the psychophysical kernels to systematically underestimate the true sensory weights.

**Figure S3.**
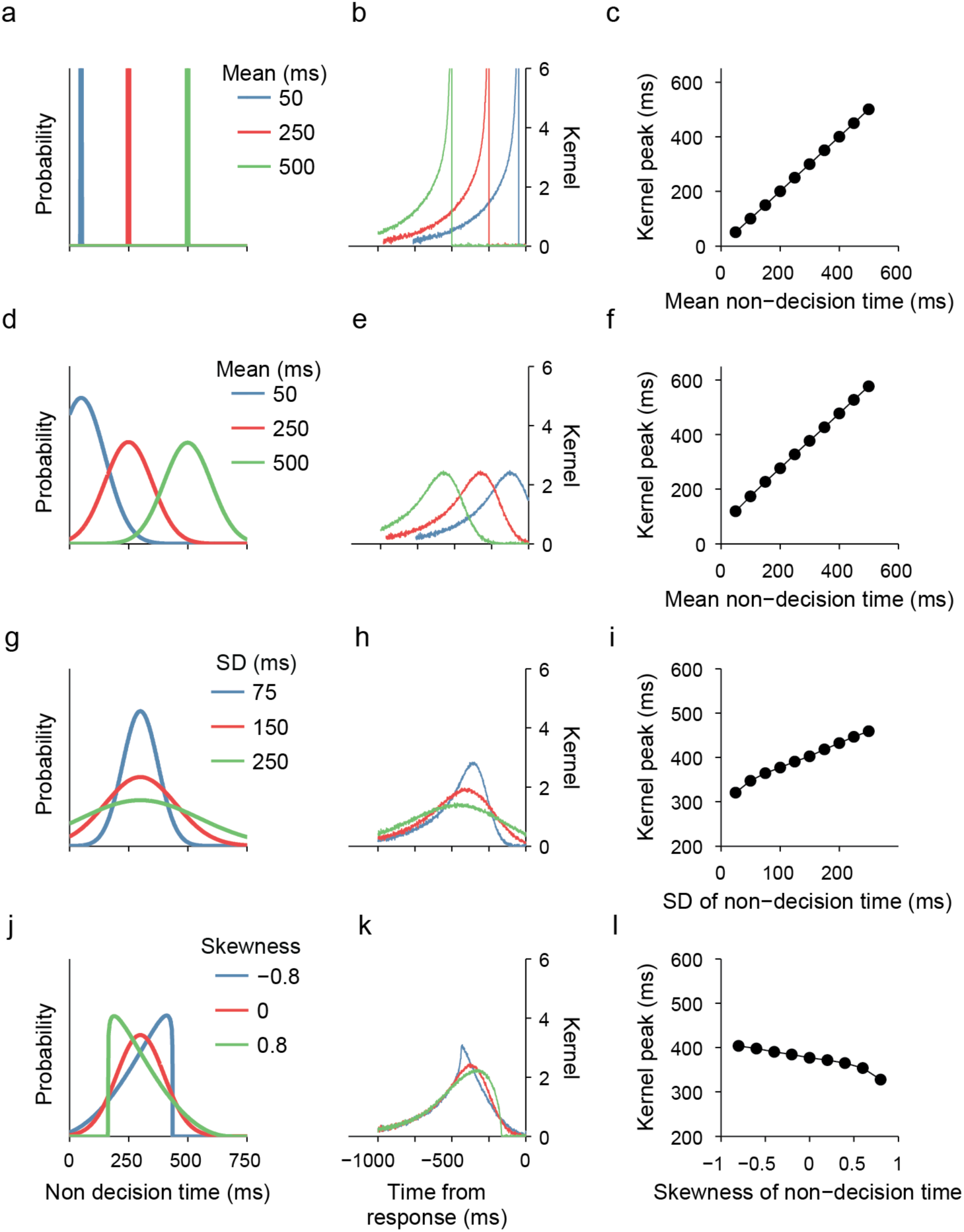
The distribution of non-decision times determines the peak time of response-aligned psychophysical kernels. Sensory and motor delays in a RT task cause the stimulus fluctuations immediately prior to the response not to bear on the choice. As a result, response-aligned kernels tend to show a peak before the response (Figs. 3-5). The time and shape of this peak are informative about the distribution of non-decision times. Here, we simulate DDMs with various distributions of non-decision time to illustrate how changes in mean, variance, and skewness of non-decision times influence the peak of psychophysical kernels. Each row illustrates three sample distributions (left column), their corresponding kernels (middle column), and systematic effects on peak time as the non-decision time distribution changes (right column). (**a-c**) When the distribution of non-decision times is a delta function (constant non-decision time), response-aligned kernels show an abrupt drop before the response. The temporal gap between this drop and response is identical to the non-decision time. (**d-f**) When non-decision time is variable (e.g., a Gaussian or truncated Gaussian distribution), changes in mean non-decision time shifts the kernel peak time, without changing its shape. Standard deviation and skewness of the distributions are set to 100 and 0, respectively. (g-i) Larger variance of non-decision time widens the kernel and shifts its peak away from the response. Mean and skewness of non-decision time are set to 300ms and 0, respectively. (**j-l**) Skewness of non-decision time creates asymmetries in the peak. More positive skewness pushes the peak toward the response. Mean and standard deviation of non-decision time are set to 300ms and 100ms, respectively. In all simulations the sensory weight function is static (*w*(*t*) = 1) and decision bound is set to 30.

**Figure S4.**
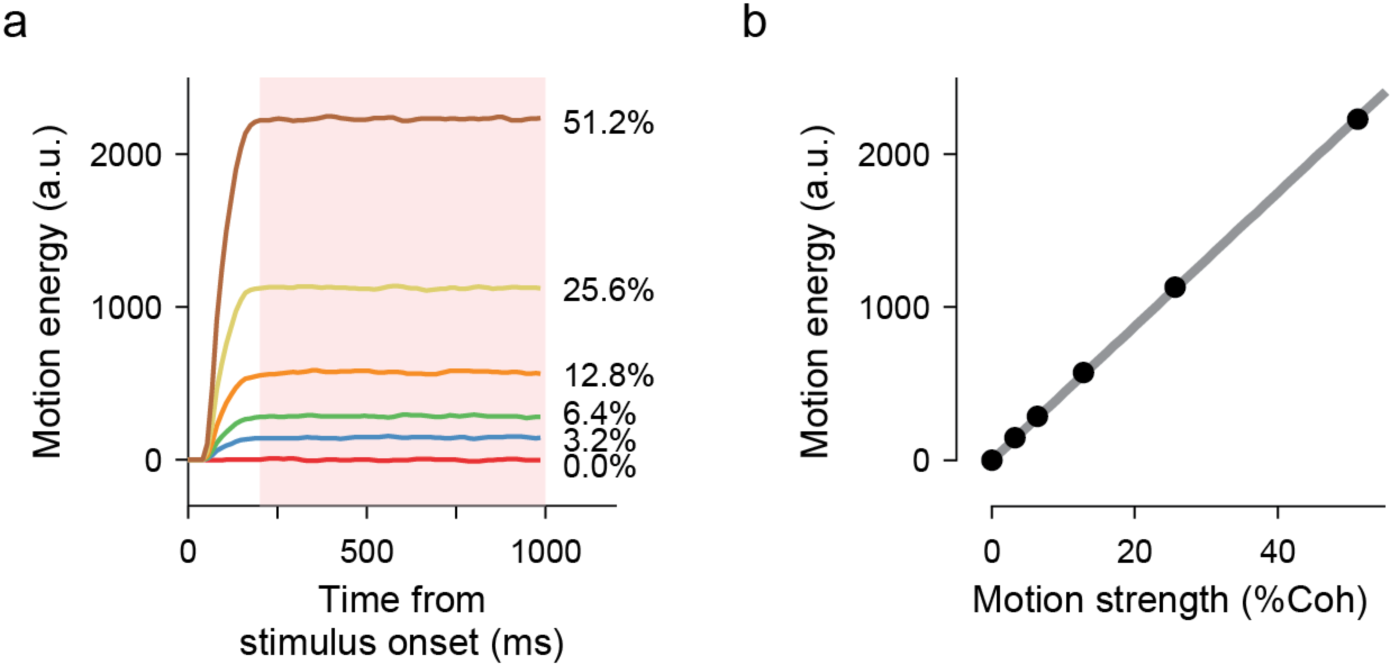
Mean motion energy is a linear function of net motion strength (coherence). (**a**) Time course of average motion energy for different coherence levels in the direction discrimination task. To calculate the scaling between motion coherence and motion energy, we computed the average motion energy 200-1000ms after stimulus onset (pink rectangle). (**b**) Average motion energy as a function of motion coherence.

**Figure S5.**
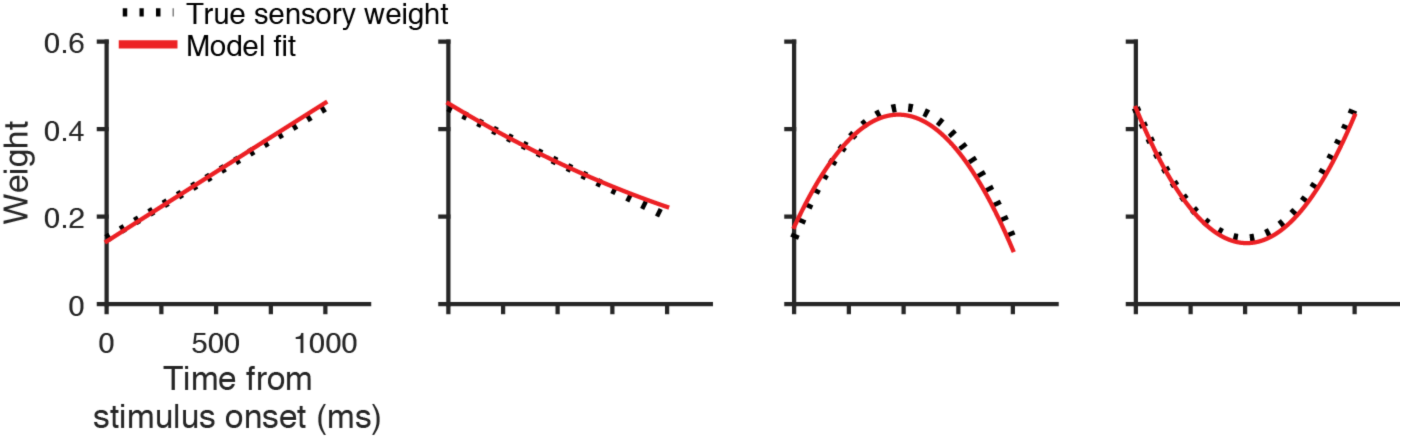
An extended DDM can recover changes of sensory weights, when such changes are present. The figure shows four simulations with different weight dynamics and the recovered weights of the model. For each panel, we simulated a direction discrimination dataset with 5000 trials. Like the real task explained in the paper, motion strength on each trial was selected randomly from a fixed set (0%, 3.2%, 6.4%, 12.8% or 25.6%). The weights in each trial could change according to a second order polynomial function. Simulated sensory evidence at each moment in a trial was a random draw from a Gaussian distribution with s.d. = 1 and mean = *w*(*τ*)*s*, where s is the motion coherence on the trial. Momentary evidence was accumulated until a positive or negative bound was reached (bound height, 30). The bound dictated the choice and time to bound was decision time. Reaction time was the sum of decision time and a random non-decision time drawn from a Gaussian distribution with mean = 300ms and s.d. = 100ms. We fit the extended DDM model with polynomial weight dynamics (Eq. 23) to the distribution of choices and RTs of the 5000 simulated trials in each panel.

**Figure S6.**
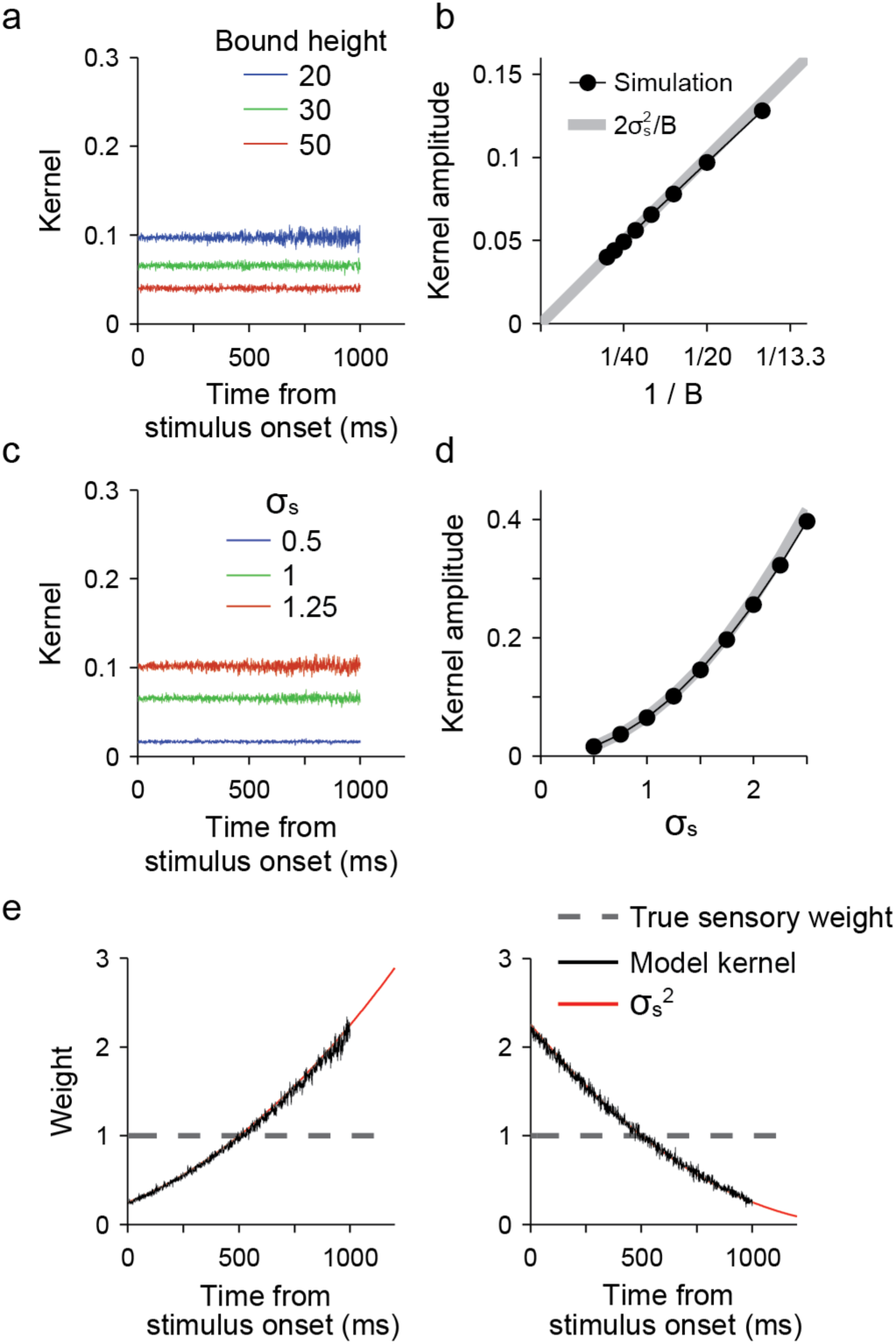
Scaling of psychophysical kernels by bound height and stimulus noise in a bounded DDM. As we prove in Methods, the psychophysical kernel of a bounded DDM without non-decision time is proportional to the sensory weight and the constant of proportionality is 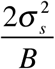 (Eq. 2). Here we confirm this relationship by simulating the model. (**a-b**) Higher decision bounds produce lower amplitudes of psychophysical kernel (a). The mean amplitude of the kernel is proportional to the inverse of decision bound (b). The gray line shows expected kernel amplitude based on Eq. 2 and the black lines and points show simulated amplitudes. 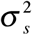 is set to 1 in these simulations. (**c-d**) Higher standard deviation of stimulus fluctuation raises the amplitude of psychophysical kernel (c). The kernel amplitude is a quadratic function of *σ_s_* (d). *B* is set to 30 for these simulations. (**e**) When stimulus noise (*σ_s_*) changes within the trial, the psychophysical kernel co-varies with it, causing deviation from the true sensory weight. Note that in Figs. 3, 6, 7, S1-2, and S7-9, we normalized the kernel based on Eq. 2 and Eq. 14 to remove the effect of bound height and stimulus variance in order to compare the kernel directly with the sensory weight.

**Figure S7.**
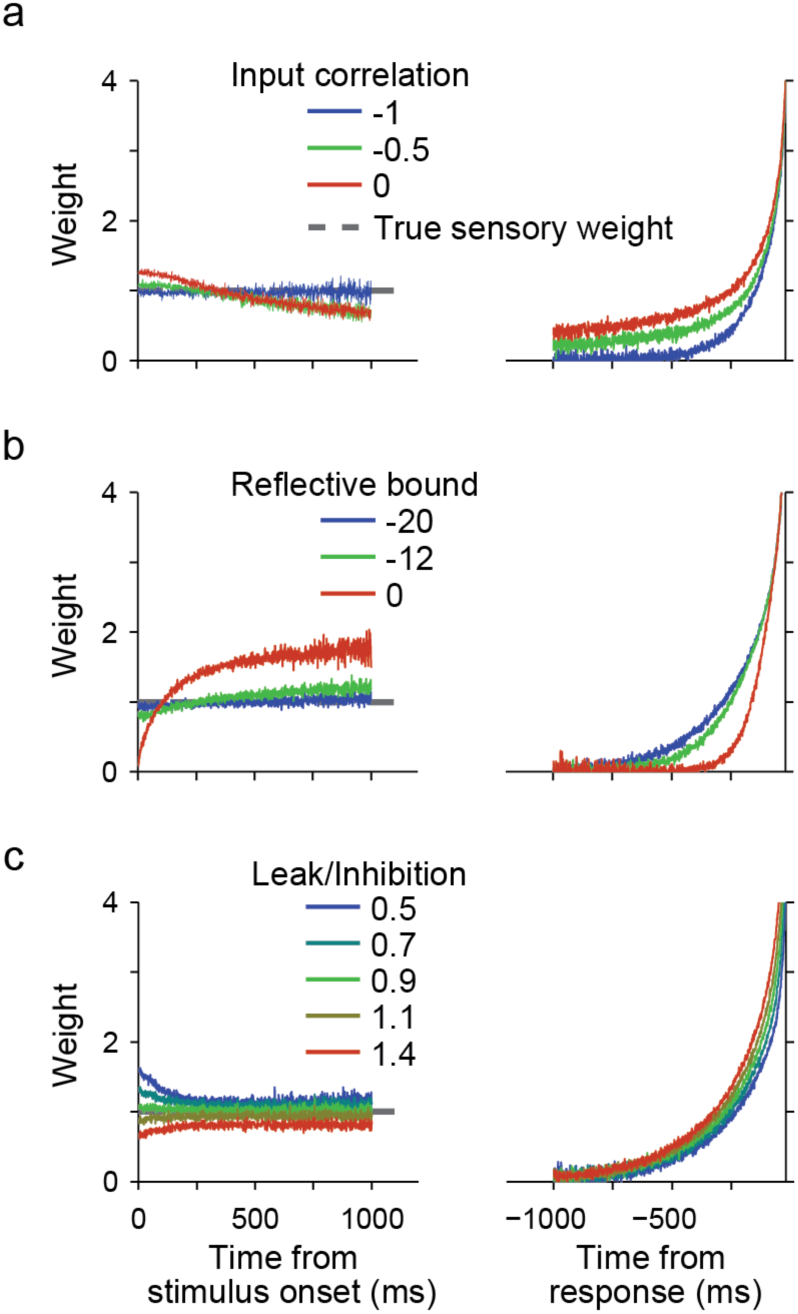
Additional examples of psychophysical kernels for various parameterizations of the competing accumulator model. To provide better intuition for the effects of model parameters in Fig. 6, we have plotted psychophysical kernels for multiple parameter values. a-d correspond to rows 2-4 in Fig. 6.

**Figure S8.**
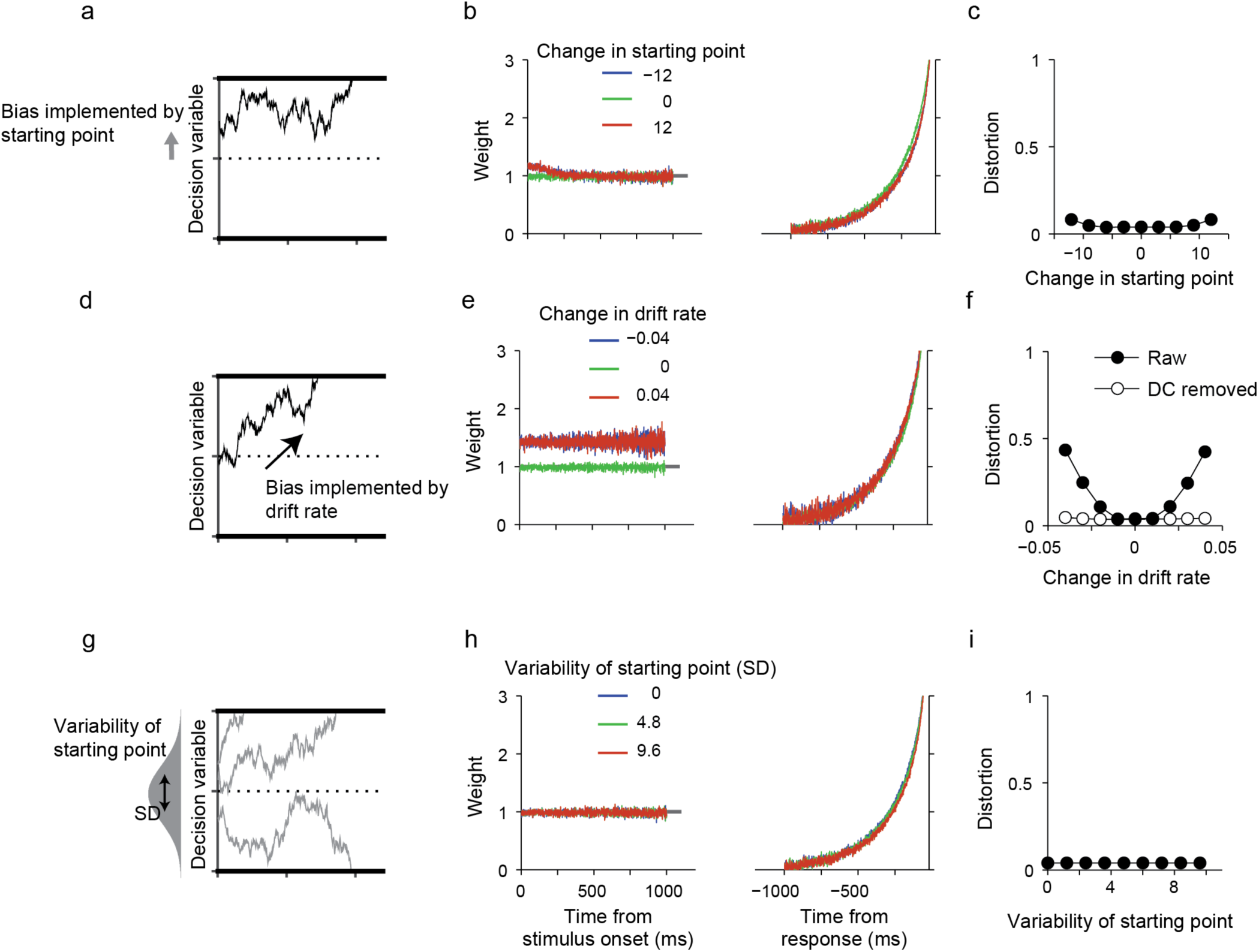
The effect of bias and variability of the starting point of the DDM on psychophysical kernels. Conventions are similar to Fig. 6, where the first column shows the model variation, the second column shows example parameterizations of the model, and the third column shows the magnitude of distortion of the psychophysical kernel as a function of the parameter of interest. Decision bound is set to 30 in all simulations. (**a-c**) When choice bias is implemented by a shift of the starting point toward a decision bound, the kernel shows a small inflation around stimulus onset, because the closer distance of the starting point to one of the bounds increases the likelihood of bound crossing due to early stimulus fluctuations. (**d-f**) When choice bias is implemented by a constant change in drift rate, the kernel shows a DC offset. (**g-i**) Trial-to-trial variability of the starting point of the DDM does not cause a systematic distortion of psychophysical kernels, if the starting point distribution is centered on zero.

**Figure S9.**
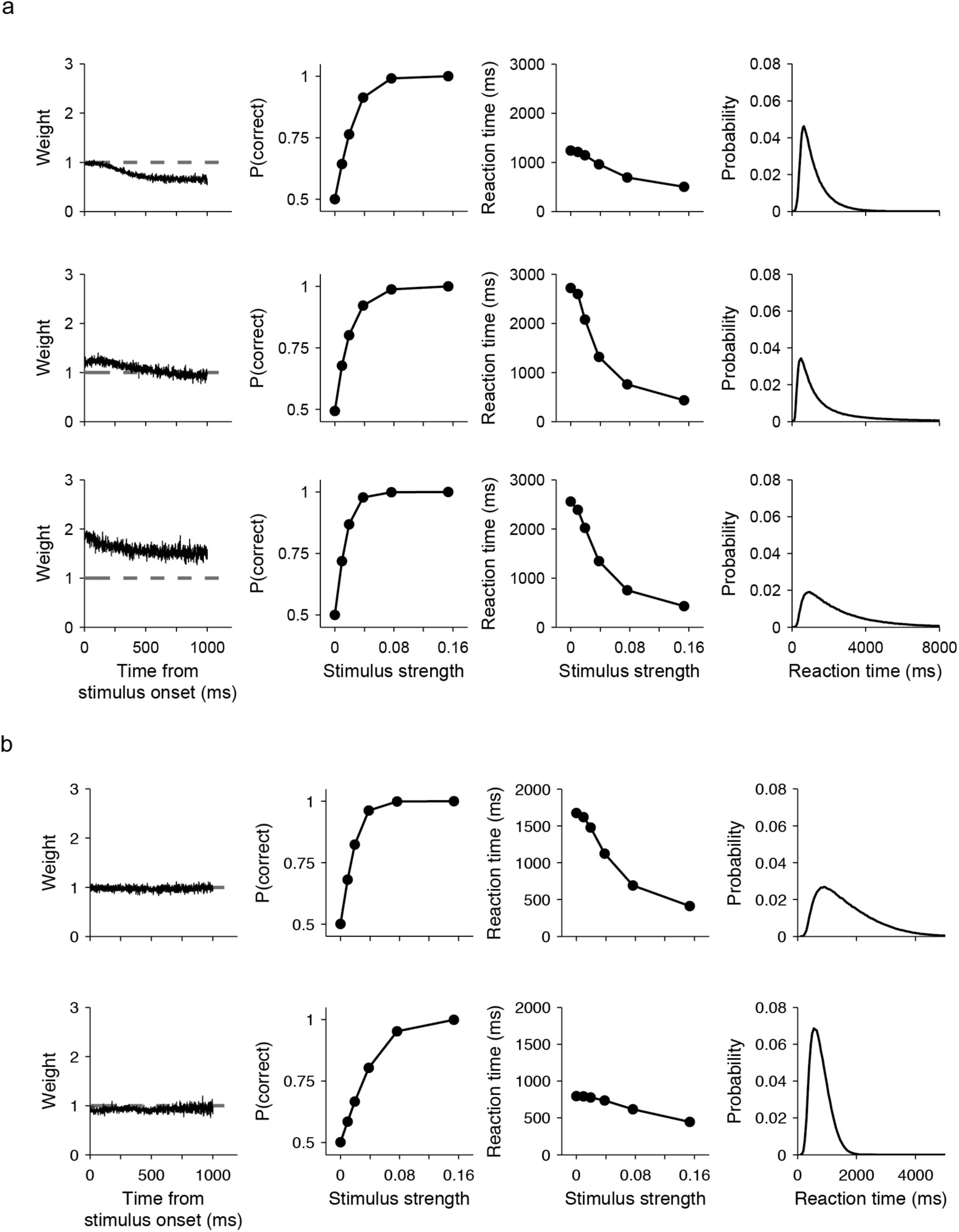
A three-pronged approach based on the shape of psychophysical kernels and the distribution of choices and RTs can distinguish different mechanisms that contribute to the decision-making process. (**a**) A diversity of mechanisms can lead to similar trends in psychophysical kernels but they usually lead to contrasting psychometric and chronometric functions, different RT distributions, and quantitative differences in the shape of kernels. The figure shows three mechanisms that cause a downward trend in stimulus-aligned kernels. The top row shows a DDM with mean non-decision time set to 300ms and s.d. of non-decision time to 100ms (B = 30). The middle row shows a competing accumulator model where the input correlation of the two accumulators is −0.1 (Leak and inhibition are set to 0; non-decision time, mean, 100ms, s.d., 33ms; B = 50). Bottom row shows a competing accumulator model where the leak to inhibition ratio is 0.8 (*L*, 0.0027; *I*, 0.0033; input correlation, −1; non-decision time, mean, 100ms, s.d., 33ms; B = 80, *v*_0_ = 30). RT distributions in the right column are for trials with stimulus strength of 0, the same trials used for making the psychophysical kernels. The parameters of the three simulations are adjusted to have a more or less similar drop in psychophysical kernels. Kernels are normalized according to Eq. 2. (**b**) A flat psychophysical kernel can emerge from a diversity of mechanisms, which often cause contrasting psychometric and chronometric functions, and different RT distributions. Top row shows a DDM with urgency (*τ*_1/2_, 5,000ms, *b*, 50, *u*_∞_, 50 in Eq. 15; non-decision time, mean, 100ms, s.d., 33ms). Bottom row show the competing accumulator model of Fig. 7d.

